# Phosphatidylinositol 3-kinase-independent synthesis of PtdIns(3,4)P_2_ by a phosphotransferase

**DOI:** 10.1101/2021.05.25.445663

**Authors:** Glenn F. W. Walpole, Jonathan Pacheco, Neha Chauhan, Yazan M. Abbas, Fernando Montaño-Rendón, Zetao Liu, Hongxian Zhu, John H. Brumell, Alexander Deiters, Gerald R.V. Hammond, Sergio Grinstein, Gregory D. Fairn

## Abstract

Despite their comparatively low abundance, phosphoinositides play a central role in membrane traffic and signalling. PtdIns(3,4,5)P_3_ and PtdIns(3,4)P_2_ are uniquely important, as they promote cell growth, survival, and migration. Pathogenic organisms have developed means to subvert phosphoinositide metabolism to promote successful infection and their survival within host organisms. We demonstrate that PtdIns(3,4)P_2_ is generated in host cells by effectors of the enteropathogenic bacteria *Salmonella* and *Shigella*. Pharmacological, gene silencing and heterologous expression experiments revealed that, remarkably, the biosynthesis of PtdIns(3,4)P_2_ occurs independently of phosphoinositide 3-kinases. Instead, we found that the *Salmonella* effector SopB, heretofore believed to be a phosphatase, generates PtdIns(3,4)P_2_ *de novo* via a phosphotransferase/phosphoisomerase mechanism. Recombinant SopB is capable of generating PtdIns(3,4)P_2_ from PtdIns(4,5)P_2_ in a cell-free system. Through a remarkable instance of convergent evolution, bacterial effectors acquired the ability to synthesize 3-phosphorylated phosphoinositides by an ATP- and kinase-independent mechanism, thereby subverting host signaling to gain entry and even provoke oncogenic transformation.

## Introduction

The distribution and metabolism of phosphatidylinositol and its polyphosphoinositide (PPIn) derivatives have profound consequences in eukaryotic cells. By directing vesicular traffic, gating ion channels, and orchestrating numerous signal transduction pathways, PPIns shape organellar identity and function (Balla, 2013; Dickson and Hille, 2019; Posor et al., 2015). The seven chemically unique PPIn species formed by combinatory phosphorylation of the inositol ring are in constant flux. Specialized kinases and phosphatases catalyze the chemical addition or removal, respectively, of phosphates at the D-3, D-4, and D-5 position of the inositol ring while phospholipase C isoforms detach the phospho-inositol ring from its glycerol backbone at the D-1 position. That numerous human pathologies are the result of gain-or loss-of-function mutations in these enzymes (Hakim et al., 2012; McCrea and De Camilli, 2009; Sasaki et al., 2009) is a testament to the central role that PPIns play in cellular homeostasis.

PtdIns(3,4,5)P_3_ and PtdIns(3,4)P_2_ are among the least abundant species in quiescent cells, constituting <0.1% of the total PPIns. However, in response to hormones, growth factors, and chemokines, phosphoinositide 3-kinases (PI3Ks) which phosphorylate the 3-hydroxyl group of the inositol ring can increase their abundance by as much as 100-fold (Dickson and Hille, 2019; Stephens et al., 1993). The PI3K family comprises 14 kinases, but a subset −class I and class II PI3Ks− are recognized in mammals as the sole pathway to generate PtdIns(3,4,5)P_3_ and PtdIns(3,4)P_2_. Class I PI3Ks activated downstream of G protein-coupled receptors (GPCRs), receptor and non-receptor tyrosine kinases (RTKs and NRTKs), or small GTPases such as Ras, phosphorylate PtdIns(4,5)P_2_ to yield PtdIns(3,4,5)P_3_ (Bilanges et al., 2019). Class II isoforms phosphorylate the 3-position of PtdIns(4)P to yield PtdIns(3,4)P_2_ in the plasma membrane (PM), but can also modify the 3-position of PtdIns to yield PtdIns(3)P in the endocytic pathway (Gulluni et al., 2019). Stereospecific interactions of the 3-phosphorylated inositol head group with cognate protein domains provide a means to amplify strong, local signals that coordinate cellular responses such as movement, mitogenesis, growth and metabolism, and autophagy via downstream effectors such as Rac, Akt, SGK, and mTOR (Hawkins and Stephens, 2016; Vanhaesebroeck et al., 2012). Of note, the inappropriate accumulation of these lipids by mutations or uncontrolled feed-forward mechanisms is a powerful anabolic signal that can drive oncogenesis (Fruman and Rommel, 2014; Lien et al., 2017).

Some pathogenic organisms have evolved means to enter eukaryotic cells by receptor-independent pathways (Pizarro-Cerdá et al., 2015; Walpole and Grinstein, 2020). The delivery of effector proteins into host cells can target central regulatory nodes of the cortical actin cytoskeleton to re-shape the plasmalemma, thereby driving bacterial internalization and the formation of vacuoles that serve as a safe intracellular niche for pathogen survival and proliferation. Remarkably, signaling associated with 3-phosphorylated PPIns has long been noted during entry of several enteropathogenic bacteria (Marcus et al., 2001; Niebuhr et al., 2002; Pendaries et al., 2006; Steele-Mortimer et al., 2000), but the originating lipids and the mechanism(s) underlying their formation have remained elusive. It has been explicitly assumed that this was a direct consequence of altered PI3K activity, but the putative kinase, whether host or bacterial, has not been identified. The precise nature of the PPIns generated has also remained uncertain due, in part, to the use of dual-specificity biosensors such as the pleckstrin-homology (PH) domain of Akt (Ebner et al., 2017; Manna et al., 2007). As 3-phosphorylated PPIns are now fertile clinical targets, the possible existence of additional biochemical means to regulate their generation and/or activity is of great interest.

In recent years, a new complement of genetically-encoded biosensors with high specificity and avidity for PtdIns(3,4)P_2_ or PtdIns(3,4,5)P_3_ have been devised and validated (Goulden et al., 2019; Liu et al., 2018). We took advantage of such improved probes, in combination with live cell spectroscopy, optogenetics, and *in vitro* reconstitution experiments, to revisit the spatiotemporal dynamics and underlying mechanism of 3-PPIns generation by enteropathogens. Our studies identified an unprecedented mechanism of 3-PPIns generation, independent of PI3Ks.

## Results

### Dynamic formation of 3-phosphorylated PPIns during pathogen entry and maturation

We initially investigated the possible formation and subcellular distribution of PtdIns(3,4)P_2_ during bacterial invasion, employing a genetically-encoded biosensor based on the carboxy-terminal PH domain of human TAPP1 (Dowler et al., 2000; Thomas et al., 2001) that was recently modified to improve its sensitivity (Goulden et al., 2019) (Figure S1A). As a primary model of enteropathogenic invasion, we exposed epithelial cells to *Salmonella enterica* (serovar Typhimurium, hereafter *Salmonella*) − a prevalent world-wide threat to humans and various zoonotic hosts (Branchu et al., 2018; LaRock et al., 2015). In the absence of serum, levels of membrane PtdIns(3,4)P_2_ in cultured epithelial cells are relatively low, resulting in a largely cytosolic distribution of the biosensor NES-EGFP-cPHx3 (hereafter cPHx3; Figure 1A, left panel) (Goulden et al., 2019). In stark contrast, the addition of invasion-competent *Salmonella* was associated with a remarkable enrichment of cPHx3 at the site of contact between bacteria and the host PM; this was followed by the generation of extensive cPHx3-labeled membrane ruffles and the internalization of bacteria into plasmalemmal-derived vacuoles (Figure 1A, right panels; Figure S1A-S1C; Supplementary Movies 1 and 2). Quantification of cPHx3 intensity in the bulk PM revealed an enrichment of the biosensor in this compartment that persisted long after pathogen entry (Figures 1B and S1E). Even though *Salmonella* entered the cells within 10 min of addition, elevated levels of cPHx3 at the PM were detected for >1 hour post-infection, despite the removal of extracellular bacteria by extensive washing and addition of antibiotics (see Methods). The phosphorylation of the sentinel kinase Akt, which recognizes 3-phosphorylation of the inositol ring, tracked temporally the increase in membrane cPHx3 (Figures S1E, and S1F).

**Fig. 1.**
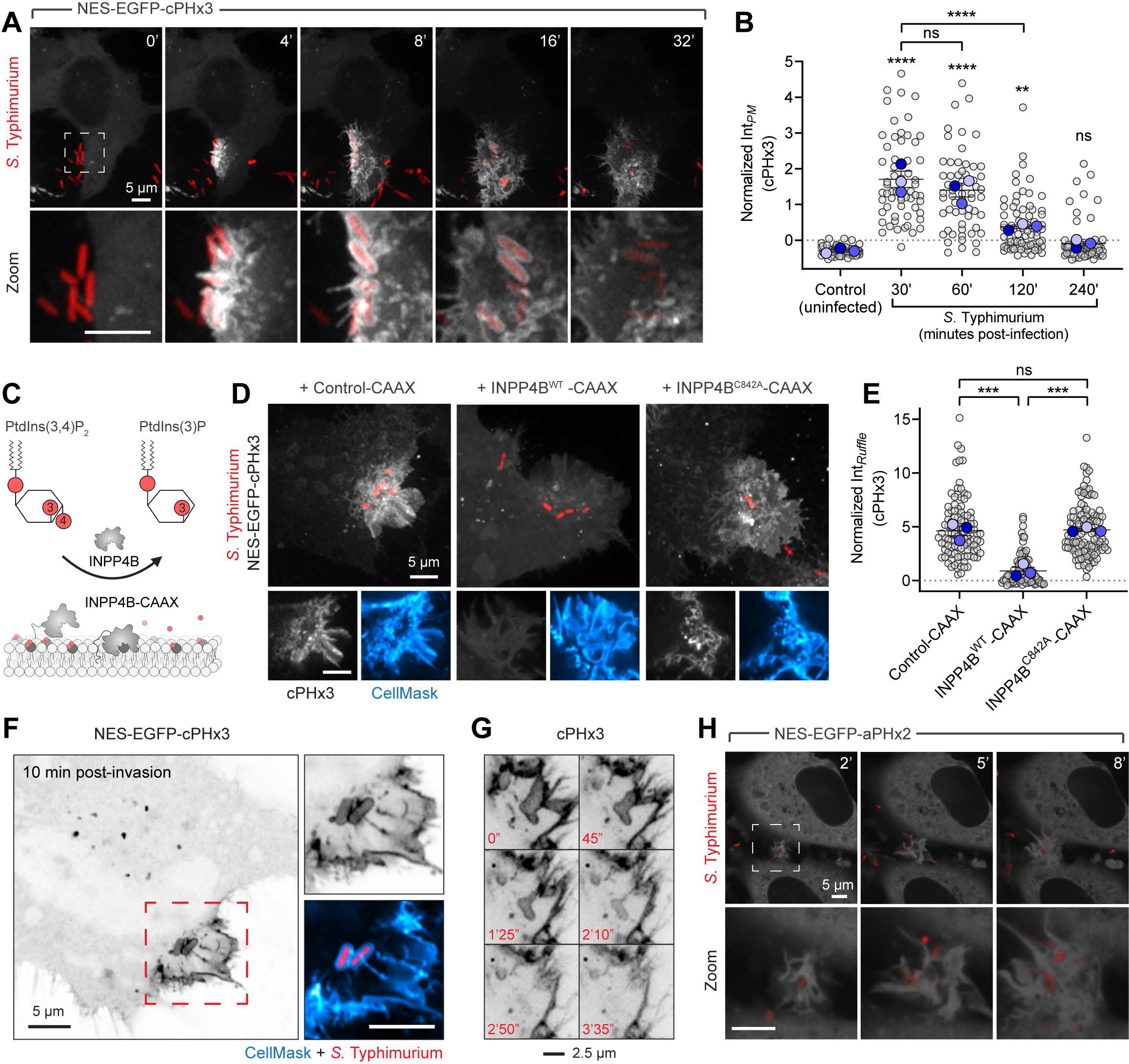
Rapid and sustained PtdIns(3,4)P_2_ synthesis during *Salmonella* entry and maturation. (A) Live cell confocal imaging of cPHx3 (gray) during invasion by RFP-expressing *Salmonella* (red). Extended focus projections (main panels) and enlargements of the hashed box region (bottom panels) are presented where 0 min indicates time of bacterial contact with the membrane. See also Fig. S1A-S1C, and Supplementary Movies 1 and 2. (B) Sustained PtdIns(3,4)P_2_ levels during infection. Cells serum-starved for 3 hours (to reduce basal PtdIns(3,4)P_2_) were exposed to *Salmonella* for 10 min. Extracellular bacteria were removed by washing, and cells returned to serum-free medium containing gentamycin (see Methods). Total PM-associated cPHx3 intensities are presented from three trials (58-75 cells per condition). Here and elsewhere, trial means ± SEM (coloured, foreground) are overlaid on cellular measurements (gray, background). Significance was assessed by one-way ANOVA of trial means with Bonferroni’s multiple comparisons test versus control (here and elsewhere ****, p < 0.0001; **, p < 0.01; ns, not significant). (C) Model of plasmalemmal PtdIns(3,4)P_2_ depletion by INPP4B-CAAX. Fusion of the D-4-specific phosphatase to the tail of HRAS preferentially targets its activity to the PM. (D and E) INPP4B impairs recruitment of the cPHx3 biosensor to *Salmonella*-induced ruffles. Normalized cPHx3 intensity in invasion ruffles was analyzed 10 min post-infection in cells co-transfected with TagBFP2-CAAX (control-CAAX, 106 ruffles), TagBFP2-INPP4B^WT^-CAAX (112 ruffles), or catalytically inactive TagBFP2-INPP4B^C842A^-CAAX (113 ruffles). Extended focus projections are presented (main), and panels below are corresponding confocal sections of the invasion ruffle. Here and elsewhere, CellMask (blue) identifies the PM. TagBFP2 fluorescence is not shown. Ordinary one-way ANOVA with Bonferroni’s post test of three trial means. Here and elsewhere (***, p < 0.001). (F and G) cPHx3 labels the invaginating region of the PM prior to fission. (F) An Airyscan micrograph is presented of cPHx3 within the invasion ruffle induced by *Salmonella* (red, 10 min post-infection). Note the continuous cPHx3 (inverted gray) labelling that coincides with a CellMask (blue, right panel)-stained membrane compartment. (G) Confocal time lapse of cPHx3 during bacterial entry and constriction of the invaginating membrane. Images are extended focus projections of a 2 μm optical slice that encompasses the bacterium (not shown). (H) Monitoring PtdIns(3,4,5)P_3_ during invasion. HeLa cells expressing aPHx2 (gray) were imaged live during invasion with *Salmonella* (red). Confocal sections are presented where 0 min marks the first indication of membrane ruffling induced by the bacteria. See also Figure S1D and Supplementary Movie 3.

To validate if cPHx3 faithfully maps a cellular pool of PtdIns(3,4)P_2_, we analyzed infection of cells overexpressing PM-targeted inositol polyphosphate 4-phosphatase type II (INPP4B), which specifically hydrolyzes the D-4 position of PtdIns(3,4)P_2_ (INPP4B-CAAX, Figure 1C) (Gewinner et al., 2009; Norris et al., 1997). The enrichment of cPHx3 within *Salmonella*-induced ruffles was largely eliminated by co-expression with active INPP4B-CAAX, but not by a point mutant (C482A) of the phosphatase lacking catalytic activity (Figures 1D and 1E). Further analyses of cPHx3 revealed that during pathogen entry, PtdIns(3,4)P_2_ became highly enriched on the forming *Salmonella*-containing vacuole, lining also the thin tubular invagination connecting it to the PM prior to scission (Figures 1A and 1F). Loss of cPHx3 on the vacuole correlated tightly with the disengagement of this membrane-bound compartment from the plasmalemma (Figures 1G and S1C). Labeling of bystander macropinocytic compartments by cPHx3 was also evident (Supplementary Movie 2).

We also monitored PtdIns(3,4,5)P_3_ distribution by utilizing tandem PH domains of ARNO (Cytohesin-2) (Cronin et al., 2004; Klarlund et al., 2000; Venkateswarlu et al., 1998). This biosensor, NES-EGFP-aPH^I303E^x2 (Goulden et al., 2019) (Figure S2A) and hereafter called aPHx2, evinced minimal enrichment in invasion ruffles (Figures 1H, S1D, S2A-S2C, Supplementary Movie 3). Similar negative results were obtained using another PtdIns(3,4,5)P_3_-specific probe comprising the PH domain of Btk (not shown). Thus, we verified that *bona fide* 3-phosphorylated PPIns are generated acutely during *Salmonella* entry; however, the predominant lipid appears to be PtdIns(3,4)P_2_ and not PtdIns(3,4,5)P_3_.

### Bacterial or heterologously expressed SopB promotes the acute synthesis of PtdIns(3,4)P_2_

Within the gastrointestinal lumen and during logarithmic growth, *Salmonella* expresses a complement of virulence factors, pre-synthesized and primed for delivery into the host cytosol for invasion. A specialized ‘molecular needle’ expressed on the bacterial surface, a type three section system, penetrates the host membrane to translocate the effector load (Galán et al., 2014). *Salmonella* outer protein B (SopB, also called SigD), one such effector, is a key virulence gene and inflammatory determinant in *Salmonella* infections (Anderson Norris et al., 1998; Galyov et al., 1997; Kum et al., 2011; Tahoun et al., 2012; Zhang et al., 2002). Remarkably, as many as half of the ≈9500 non-redundant phosphorylation events that arise in response to infection are SopB-dependent (Rogers et al., 2011). The functions of SopB have been tightly linked to its enzymatic activity, aforethought to be a phosphatase which acts on inositol phosphates and phosphoinositide lipids of the host cell (Anderson Norris et al., 1998; Feng et al., 2001; Marcus et al., 2001).

We tested the requirement of SopB for the generation of PtdIns(3,4)P_2_ by comparing host responses to wild-type *Salmonella* or an isogenic strain devoid of this effector (Δ*sopB*). As shown in Figures 2A and 2B, invasion of epithelial cells by the wild-type strain consistently induced the recruitment of cPHx3 to the invasion site; however, invasion ruffles induced by bacteria lacking SopB were completely devoid of this enrichment. Further, the overall increase in plasmalemmal PtdIns(3,4)P_2_ following invasion (Figure 1B) and the elevated phosphorylation of Akt also required the effector SopB (Figures S1E and S1F); the latter observation is consistent with earlier findings (Steele-Mortimer et al., 2000). Similarly, the marginal enrichment of PtdIns(3,4,5)P_3_ detected by the sensor aPHx2 was negligible during invasion by Δ*sopB* bacteria (Figures S2A-S2C).

**Fig. 2.**
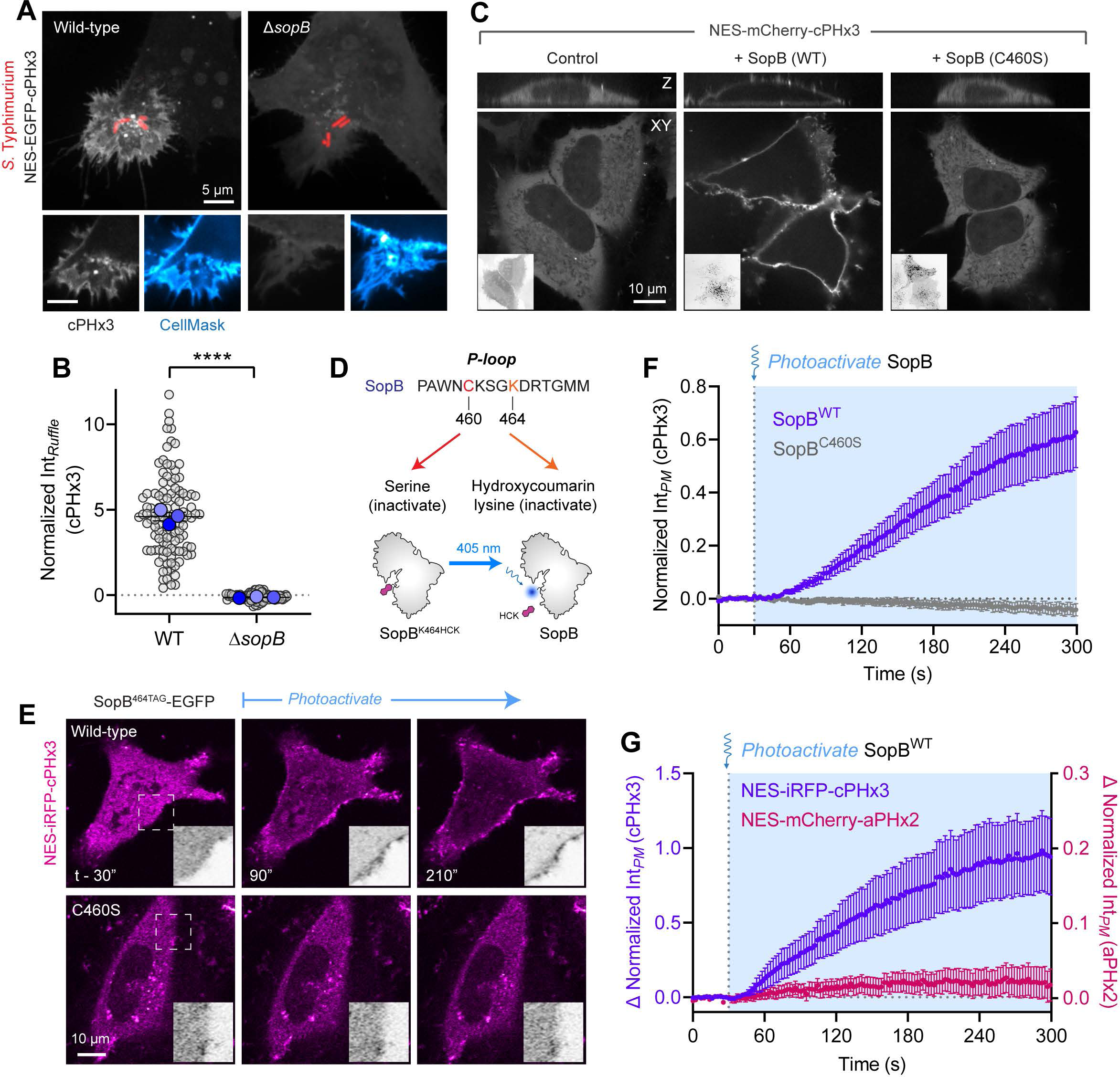
SopB is necessary and sufficient for PtdIns(3,4)P_2_ biosynthesis in mammalian cells. (A) Requirement of SopB for PtdIns(3,4)P_2_ generation. Cells expressing cPHx3 were infected for 10 min with wild-type or Δ*sopB Salmonella* expressing RFP (red) prior to staining the PM (blue). Extended focus projections (main) and corresponding confocal sections of the invasion ruffle (bottom panels) are presented. (B) Normalized membrane cPHx3 intensity was quantified in 3 independent trials (WT, 113 ruffles; Δ*sopB*, 87 ruffles). Unpaired two-tailed t-test. (C) Heterologous expression of SopB suffices to generate PtdIns(3,4)P_2_. HeLa cells were co-transfected as indicated prior to imaging live and the distribution of mCherry-labeled cPHx3 is shown. Representative XY and Z sections are presented from 3-5 independent trials (N) per construct (Control (EGFP only), 122 cells, N=5; SopB^WT^-EGFP, 210 cells, N=5; SopB^C460S^-EGFP, 107 cells, N=3). Insets depict the GFP signal (inverted grey scale). (D) Optogenetic activation of SopB. Hydroxycoumarin lysine (HCK) is incorporated at position 464 by TAG codon mutagenesis to block the active site. Cells are transfected with SopB^464TAG^ and plasmids that encode an Amber stop codon (UAG)-recognizing tRNA and a tRNA synthase that incorporates HCK. Illumination by 405 nm light photolyzes the hydroxycoumarin to yield wild-type SopB. A dual SopB C460S-K464TAG negative (enzymatically-inactive) control was also generated. (E) Photoactivation of SopB induces acute PtdIns(3,4)P_2_ formation. Representative time-lapse of cells expressing cPHx3 (magenta) pre- and post-photoactivation of SopB^WT-464TAG^ or SopB^C460S-464TAG^. Insets are inverted grayscale images of an expanded region of the PM marked by the hashed box in the left-most frame. See also Supplementary Movie 4. (F) Quantification of experiments like those in (E) from three independent trials (SopB^WT^, 119 cells; SopB^C460S^, 50 cells). Significance of the difference between the areas under curves, assessed by two-tailed Mann-Whitney was p < 0.0001 (not shown). Here and elsewhere, cyan-shaded areas represent 405-nm-illuminated timepoints and data are means ± SEM of individual cells. See also Supplementary Fig. 2. (G) Scarcity of PtdIns(3,4,5)P_3_ during photoactivation of SopB. Quantification of the PM intensity of NES-iRFP-cPHx3 (left y-axis, purple) and NES-mCherry-aPHx2 (right y-axis, pink) following the activation of SopB^WT^ (3 trials, 29 cells).

The possibility that SopB acts in concert with other virulence factors to initiate this response was examined next. To exclude the possible contribution of other bacterial effectors translocated during infection we transfected epithelial cells with a plasmid encoding SopB. Heterologous expression of SopB alone led to a marked translocation of cPHx3 to the PM of the transfected cells (Figure 2C), implying that PtdIns(3,4)P_2_ synthesis occurs independently of other bacterial effectors. SopB shares homology with the Cys-X_5_-Arg consensus sequence of the active site of the mammalian D-4 phosphatases INPP4A/4B and the D-3 phosphatase PTEN (Anderson Norris et al., 1998). Mutation of the homologous cysteine in SopB to serine (C460S), reported to block its phosphatase activity (Anderson Norris et al., 1998; Marcus et al., 2001), also completely blocked its effect on cPHx3 translocation (Figure 2C, right). These results imply that the catalytic activity of SopB is necessary to mediate the burst of PtdIns(3,4)P_2_ during *Salmonella* invasion.

Unlike its acute response revealed during bacterial invasion, manifestation of the effects of SopB on PtdIns(3,4)P_2_ in the case of heterologous SopB expression require hours, raising the possibility that they may arise indirectly, through slow intermediate events. To investigate this possibility we turned to acute optogenetic activation of the enzyme (Courtney and Deiters, 2019; Luo et al., 2014). We selected K464 of SopB, whose codon was mutated to UAG and co-expressed with a plasmid to facilitate the incorporation of the unnatural, caged amino acid, hydroxycoumarin lysine (HCK) during protein translation (Figure 2D, see Methods). We predicted that the active site would be occluded –and thus non-functional– by the bulky hydroxycoumarin derivative. As predicted, the transfection of SopB^464TAG^ did not support the generation of PtdIns(3,4)P_2_ (Figure 2E, top left panel). However, light-induced photolysis of the HCK group by 405-nm light illumination restored the K464 residue (Figure 2D), leading to rapid (< 1 min) SopB-mediated formation of PtdIns(3,4)P_2_ in the PM (Figures 2E and 2F, Supplementary Movie 4). Importantly, the increase in PtdIns(3,4)P_2_ was not observed when the C460S mutation was introduced into SopB^464TAG^ (Figures 2E and 2F), ruling out a spurious effect of light exposure. Using this approach we compared the dynamics of PtdIns(3,4)P_2_ and PtdIns(3,4,5)P_3_ generation during the acute photo-activation of heterologous SopB in live cells. As in the previous experiments we observed minimal generation of PtdIns(3,4,5)P_3_ in cells exhibiting robust PtdIns(3,4)P_2_ responses (Figures 2G and S2F; note expanded scale of aPHx2 plot); the marginal translocation of aPHx2 was similar in cells expressing wild-type or C460S mutant SopB (Figures S2D-S2G). Thus, SopB is both necessary and sufficient to generate a rapid and robust PtdIns(3,4)P_2_ response in human cells.

### Phosphoinositide 3-kinases are not required for SopB-driven PtdIns(3,4)P_2_ formation

Hydrolysis of the predominant cellular phosphoinositides PtdIns(4)P and PtdIns(4,5)P_2_ (comprising ≈1-2% of the lipids in the PM) by a phosphatase cannot directly account for the observed PtdIns(3,4)P_2_ synthesis. Instead, we entertained the possibility that SopB stimulates the activity of a host PI3K, which feature prominently in endocytic pathways (Marat and Haucke, 2016; Walpole and Grinstein, 2020). In response to the activation of cell surface GPCRs, RTKs and NRTKs, and specific small GTPases such as Ras, PtdIns(4,5)P_2_ is phosphorylated to PtdIns(3,4,5)P_3_ by class I PI3Ks. PtdIns(3,4,5)P_3_ can be rapidly converted to PtdIns(3,4)P_2_ by subsequent dephosphorylation by several capable 5-phosphatases (Balla, 2013; Sasaki et al., 2009) (Figure 3A). Indeed, the constitutive recruitment of a class I PI3K to the PM (p110α-CAAX) triggered a parallel increase in PtdIns(3,4)P_2_ levels (Figures 3B and 3C).

**Fig. 3.**
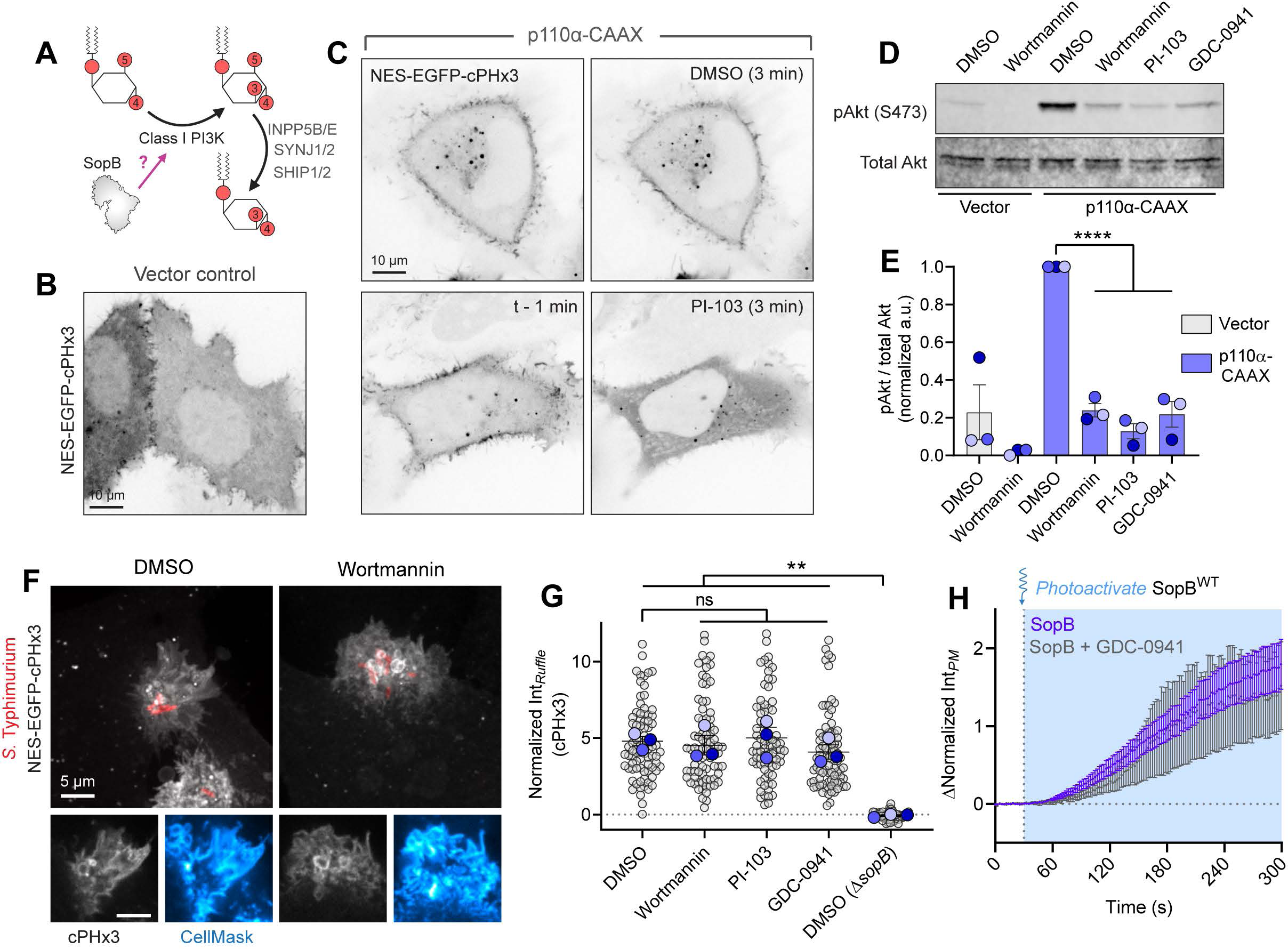
Class I PI3Ks are not required for PtdIns(3,4)P_2_ stimulated by SopB. (A) Model of a PI3K-mediated, indirect PtdIns(3,4)P_2_ synthesis in mammalian cells and potential site of action of SopB. SopB could conceivably stimulate class I PI3Ks and the resulting PtdIns(3,4,5)P_3_ could subsequently be dephosphorylated to PtdIns(3,4)P_2_ by multiple 5-phosphatases. (B and C) Indirect generation of PtdIns(3,4)P_2_ by class I PI3Ks is rapidly terminated by PI3K inhibitors. Cells expressing cPHx3 (inverted gray) were co-transfected with vector control (B) or with the PM-targeted catalytic subunit of class IA PI3K (C). Cells expressing p110α-CAAX were subsequently treated with DMSO (vehicle; top row) or (500 nM; bottom row). Images were acquired immediately before (left panels) or 3 min after addition of DMSO or PI-103; the latter caused rapid release of cPHx3 from the PM. Similar results were obtained with 100 nM wortmannin (not shown). Micrographs shown are representative of three experiments. (D and E) Reversal of class IA PI3K-induced AKT phosphorylation by a panel of PI3K inhibitors. HeLa cells expressing vector control or p110α-CAAX were treated for 20 min with DMSO, wortmannin (100 nM), PI-103 (500 nM), or GDC-0941 (500 nM), as indicated, prior to collection of cell lysates and immunoblotting using phospho-AKT (S473) and pan AKT antibodies. The immunoblot presented is representative of three trials. (E) Densitometric quantification of 3 blots like that in D, normalized to the DMSO + p110α-CAAX condition. Data are means ± SEM and significance was calculated by one-way ANOVA with Bonferroni’s multiple comparisons test versus the DMSO control. See also Supplementary Fig. 3F. (F) Cells expressing cPHx3 (gray) were pre-treated as in (D) for 20 min prior to exposure to wildtype *Salmonella* (RFP, red). A representative extended focus projection of a DMSO-(control) or Wortmannin (100 nM)-treated cell is presented (main) with corresponding confocal sections of the invasion ruffle (bottom panels). (G) Quantification of cPHx3 intensity in the invasion ruffle from experiments like that in (F) following pre-treatment with the pan-PI3K inhibitors wortmannin or PI-103, or the class I inhibitor GDC-0941. The experiment was repeated with similar results 3 times quantifying the following number of ruffles per condition: DMSO, 81; wortmannin, 80; PI-103, 76; GDC-0941, 85; DMSO with Δ*sopB* mutant, 58. p-values derived from one-way ANOVA with Bonferroni’s multiple comparisons test versus DMSO conditions (**, p < 0.01). (H) PtdIns(3,4)P_2_ generation by heterologous SopB persisted despite PI3K inhibition. HeLa cells were left untreated or treated with GCD-0941 (250 nM) for 30 min prior to photoactivation (t=30 s) of SopB^WT-464TAG^. Data are means ± SEM of individual cell measurements (control, 40 cells; GDC-0941, 7 cells).

We employed several potent pan-PI3K inhibitors to probe the role of this pathway: wortmannin, a fungal metabolite which irreversibly modifies the active site of multiple PI3K isoforms (Arcaro and Wymann, 1993); PI-103, a synthetic multi-target PI3K and mTOR inhibitor (Knight et al., 2006); and Pictilisib (GDC-0941) which exhibits selectivity towards class I PI3Ks (Folkes et al., 2008). As shown in Figures 3D, 3E and S3F, the addition of a nanomolar concentrations of these compounds to cells expressing p110α-CAAX greatly reduced phosphorylation of Akt and led to the complete and rapid (within 3 min) release of cPHx3 from the PM (Figure 3C), validating their potency towards PI3Ks. Remarkably, despite their obvious efficacy towards PI3Ks, pre-incubation of cells with any of the above inhibitors failed to reduce PtdIns(3,4)P_2_ biogenesis at the site of *Salmonella* invasion (Figures 3F and 3G) or to block synthesis of PM PtdIns(3,4)P_2_ following optogenetic activation of SopB (Figure 3H).

As an alternative to SopB stimulating a class I PI3K, we tested the role of class II PI3Ks which can directly phosphorylate PtdIns(4)P to generate PtdIns(3,4)P_2_ (Figure 4A). It is noteworthy that of the class II isoforms, only PI3K-C2α is (somewhat) refractory to classical inhibitors of PI3Ks (Domin et al., 1997; Knight et al., 2006; Virbasius et al., 1996) and may have resisted inhibition in our initial pharmacological screen (Figure 3). To test the role of PI3K-C2α in mediating the formation of PtdIns(3,4)P_2_ by SopB, two independently targeting siRNA sequences against PIK3C2A (encoding PI3K-C2α) were introduced into the cells in conjunction with the biosensor cPHx3. Despite depletion of 85-95% of the class II PI3K, determined at the protein level by immunoblotting (Figures 4B, 4C, and S4A), the robust recruitment of cPHx3 to *Salmonella*-induced ruffles persisted (Figures 4D and 4E). Moreover, endogenous PI3K-C2α showed no enrichment at invasion ruffles (Figures S4B and S4C). Together, these observations raised the possibility that classically defined biosynthetic pathways are not responsible for the PtdIns(3,4)P_2_ synthesis induced by SopB.

**Fig. 4.**
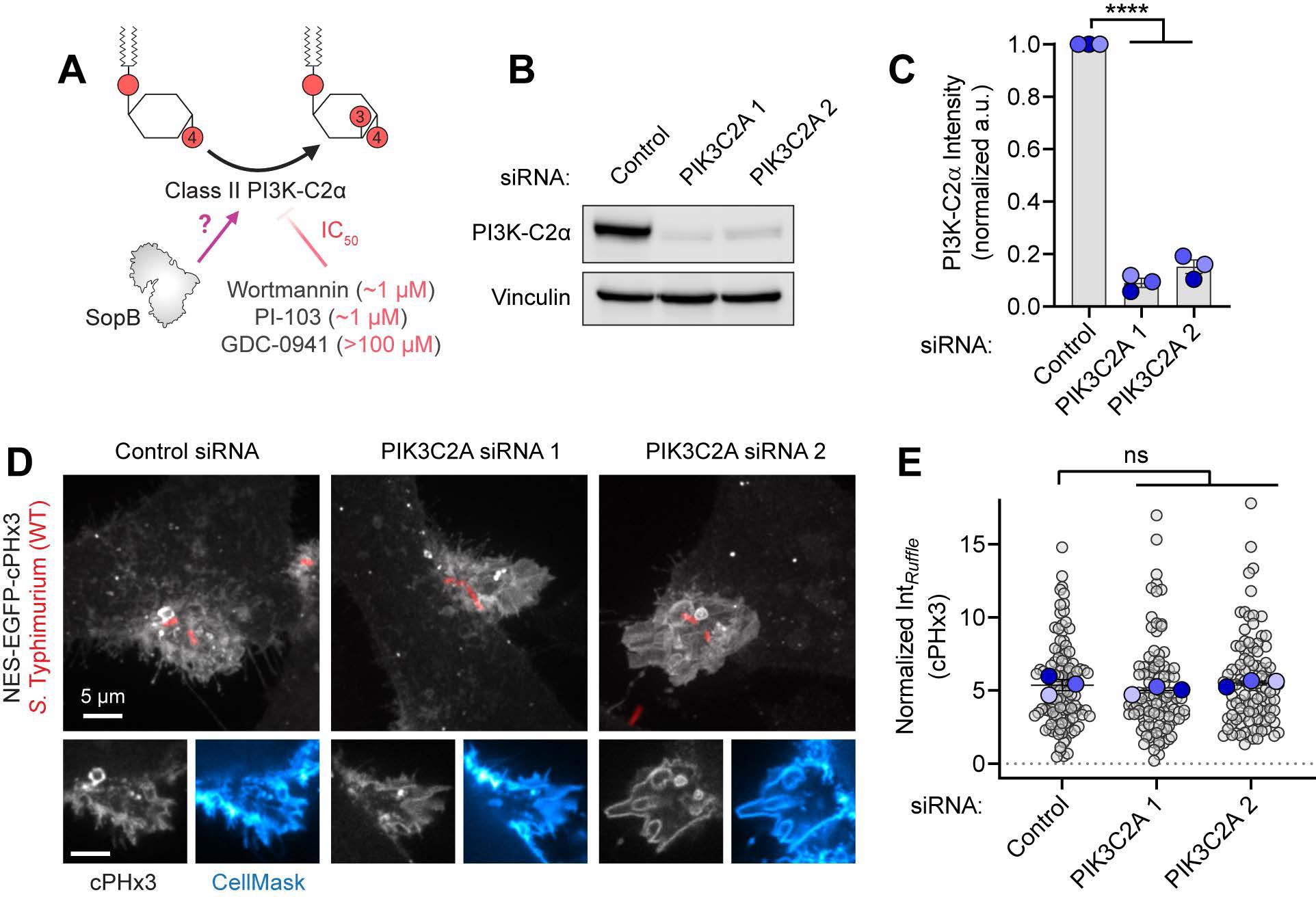
Class II PI3K-C2α is not required for PtdIns(3,4)P_2_ synthesis during *Salmonella* entry. (A) Potential SopB-activated pathway for PtdIns(3,4)P_2_ generation. The class II PI3K-C2α is largely insensitive to the phosphoinositide-3 kinase inhibitors wortmannin, PI-103, and GDC-0941 (IC_50_ values of 1 μM, 1 μM, and >100 μM respectively). (B) RNAi-mediated depletion of PI3K-C2α. Cells were treated with non-targeting or with two independent siRNAs targeting PIK3C2A and used to prepare lysates for immunoblotting against PI3K-C2α. Vinculin was utilized as a similar MW loading control. Blot is representative of 3 similar experiments. (C) Densitometric estimation of the fraction of PI3K-C2α remaining in the cells after RNAi treatment, normalized to vinculin; data from 3 independent trials. One-way ANOVA with Bonferroni’s multiple comparison test. See also Supplementary Fig. 4. (D) Cells treated with non-targeting or with two independent siRNAs targeting PIK3C2A were transfected with cPHx3 (gray) and infected with wild-type *S.* Typhimurium (red) for 10 min before staining the PM (blue). Representative extended focus projections (main) and corresponding confocal sections of the invasion ruffle (bottom) are presented. (E) Quantification of cPHx3 intensity at the invasion ruffle in 3 trials in the indicated number of ruffles (control, 117; PIK3C2A 1, 105; PIK3C2A 2, 107). One-way ANOVA with Bonferroni’s multiple comparison test (ns, not significant).

### IpgD from Shigella also generates PtdIns(3,4)P_2_ independently of phosphoinositide 3-kinases

Based on sequence homology (Figure S3A), we hypothesized that SopB and IpgD –a *Shigella flexneri* effector– share analogous enzymatic activities. Using an infection-free system, we monitored cPHx3 localization when heterologously co-expressed with IpgD. Indeed, expression of wild-type IpgD –but not IpgD with cysteine 439 mutated to serine– led to a robust relocalization of cPHx3 to the PM, with associated depletion of the biosensor from the cytosol (Figure S3D). The role of PI3Ks as possible mediators of the effect of IpgD was interrogated by addition of PI-103 and wortmannin. Quantifying cPHx3 intensity in the bulk PM revealed that, as in the case of SopB, IpgD-mediated generation of PtdIns(3,4)P_2_ was unaffected by PI3K inhibition (Figures S3D and S3E). Thus, *Shigella*’s IpgD and *Salmonella*’s SopB likely share their mode of action, apparently employing an unrecognized pathway for PtdIns(3,4)P_2_ generation in host cells.

### SopB activity is observed in Saccharomyces cerevisiae, an organism devoid of Class I and II PI3Ks

The inhibitor studies and silencing of PI3K-C2α strongly suggest that the class I and II kinases are not required for SopB-mediated PtdIns(3,4)P_2_ production. To assess this conclusion more definitively we examined the effect of SopB in *Saccharomyces cerevisiae*, an organism lacking class I and class II PI3Ks (Schu et al., 1993; Vanhaesebroeck et al., 2010). SopB was expressed acutely under the control of a galactose-inducible promoter because we found that its prolonged expression was deleterious to the yeast. When expressed in otherwise untreated *S. cerevisiae* the cPHx3 probe was entirely cytosolic (Figure 5A, left panel), consistent with their inability to synthesize PtdIns(3,4)P_2_ endogenously. Remarkably, however, heterologous expression of SopB in the yeast yielded a robust translocation of cPHx3 to the PM and a concomitant decrease in cytosolic fluorescence (Figure 5A). Importantly, the cysteine residue at position 460 of SopB was strictly required for the translocation of cPHx3 to the PM of *S. cerevisiae* (Figures 5A and 5B).

**Fig. 5.**
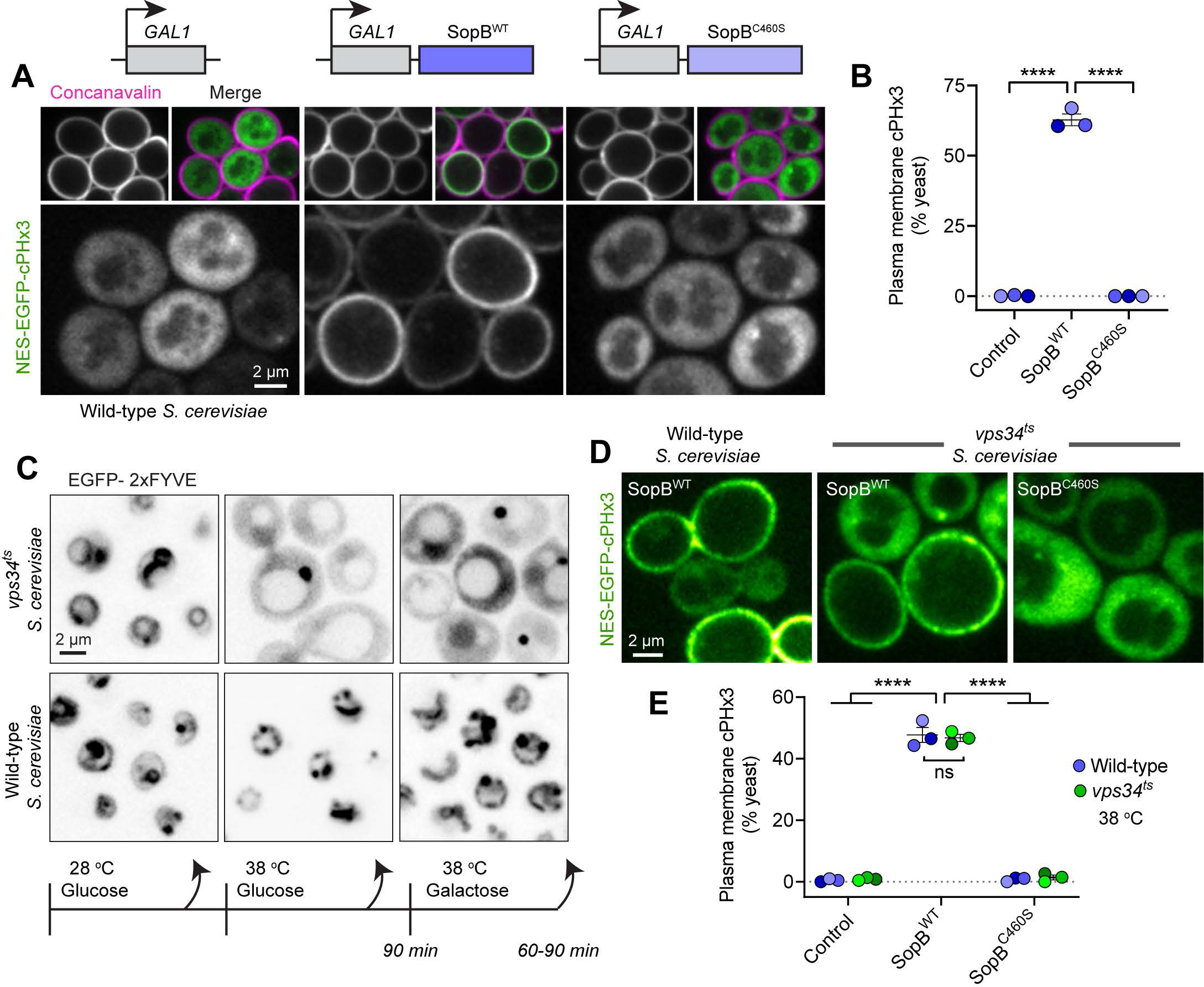
PtdIns(3,4)P_2_ generation in *S. cerevisiae* that are devoid of PI3K activity. (A) PM PtdIns(3,4)P_2_ synthesis in *S. cerevisiae*. Galactose-inducible empty vector (control), SopB^WT^, or SopB^C460S^ were induced for 2 hours in yeast that expressed NES-EGFP-cPHx3 (green in merge). Concanavalin staining (magenta) demarcates the cell wall and trypan blue was added to exclude non-viable yeast (not shown). (B) Yeast from (A) were scored for plasmalemmal cPHx3 localization across 3 trials (control, 929 cells; SopB^WT^, 929 cells; SopB^C460S^, 1116 cells). Means of individual experiments are shown; significance calculated by one-way ANOVA with Bonferroni’s multiple comparisons test. (C) PtdIns(3)P loss in *vps34^ts^* mutant *S. cerevisiae*. Cells expressing EGFP-2xFYVE (inverted gray) were maintained at 28-30°C (permissive) during growth or shifted where indicated to 38 °C (non-permissive) for 90 min before switching carbon source and continued growth at 38 °C. (D) PtdIns(3,4)P_2_ synthesis by SopB persisted despite loss of Vps34 activity. Yeast expressing galactose-inducible plasmids (see A) were shifted to a non-permissive temperature as in (C) before imaging. Non-viable cells were excluded using Trypan blue (not shown). (E) Quantification of cPHx3 translocation to the PM from experiments like (D). Control (WT, 455 cells; *ts*, 510 cells), SopB^WT^ (WT, 597 cells; *ts*, 475 cells), SopB^C460S^ (WT, 518 cells; *ts*, 331 cells). Significance calculated by two-way ANOVA with Bonferroni’s multiple comparisons test.

Despite lacking class I and class II PI3Ks, it was conceivable that other yeast PI3Ks might be involved in the SopB-induced generation of PtdIns(3,4)P_2_ via a non-canonical pathway. Class III kinase (Vps34) is the sole PI3K expressed in *S. cerevisiae*. Deletion of *VPS34* and the resultant loss of its enzymatic product, PtdIns(3)P, severely compromise endo-vacuolar protein sorting and hence the growth and survival of the yeast (Herman and Emr, 1990; Schu et al., 1993). To analyze the involvement of the kinase we therefore turned to a validated temperature-sensitive mutant (*vps34^ts^*) that exhibits acute loss of endo-vacuolar PtdIns(3)P only at non-permissive temperatures, which we confirmed by monitoring the distribution of a probe consisting of tandem FYVE domains that recognizes PtdIns(3)P (Stack et al., 1995) (Figure 5C). SopB expression was induced 90 min after shifting to a non-permissive temperature in the *vps34^ts^* strain, before monitoring cPHx3 localization. Note that the conditions required for expression of SopB, i.e. addition of galactose, did not restore the formation of PtdIns(3)P. Despite the loss of Vps34 activity and cellular PtdIns(3)P, expression of wild-type –but not of C460S– SopB induced translocation of cPHx3 to the PM (Figures 5D and 5E). We concluded that neither Vps34 nor PtdIns(3)P were involved in the generation of PtdIns(3,4)P_2_.

### SopB requires PtdIns(4,5)P_2_ to generate PtdIns(3,4)P_2_

Jointly, the preceding results appear to rule out the involvement of known host PI3Ks in the generation of PtdIns(3,4)P_2_ induced by SopB; a distinct biosynthetic pathway must therefore be invoked. *In vitro*, SopB can function as a rather promiscuous phosphoinositide and inositol polyphosphate phosphatase (Anderson Norris et al., 1998; Marcus et al., 2001), and causes the disappearance of PtdIns(4,5)P_2_ from the base of the invasion ruffle *in vivo* (Terebiznik et al., 2002). These effects depend on cysteine 460, as they are obliterated by substitution to serine (C460S mutation). It is noteworthy that the same residue is also essential for the formation of PtdIns(3,4)P_2_ by SopB in mammalian and yeast cells (Figures 2 and 5). On this basis we hypothesized that, rather than activating host kinases, SopB directly causes phosphorylation of the D-3 position of the inositol ring through rearrangement of the phosphate groups of pre-existing cellular lipids.

To test this hypothesis we analyzed the fate of PPIns regio-isomers in the membrane during the course of SopB-induced generation of PtdIns(3,4)P_2_. To monitor PtdIns(4)P levels we utilized a biosensor based on tandem P4M domains from the *Legionella pneumophila* effector SidM (Hammond et al., 2014; Tóth et al., 2016). Live cell-imaging revealed that PtdIns(4)P was abundant in the resting PM as well as in *Salmonella-*induced ruffles, and could hence potentially serve as a substrate for the formation of PtdIns(3,4)P_2_ (Figure 6A). To directly test the possible involvement of PtdIns(4)P in the formation of PtdIns(3,4)P_2_, we simultaneously monitored the distribution of the P4M and cPHx3 sensors tagged with different fluorophores during optogenetic activation of SopB. Under these conditions, plasmalemmal PtdIns(4)P was largely unperturbed despite robust PtdIns(3,4)P_2_ generation in the same cells (Figures 6B and 6C). The small, statistically insignificant decrease in PtdIns(4)P observed following SopB photo-decaging was also observed when using C460S SopB, implying that it was unrelated to the generation of PtdIns(3,4)P_2_ (Figures 6D and 6E). Thus, we found no evidence that PtdIns(4)P was directly involved in the process.

**Fig. 6.**
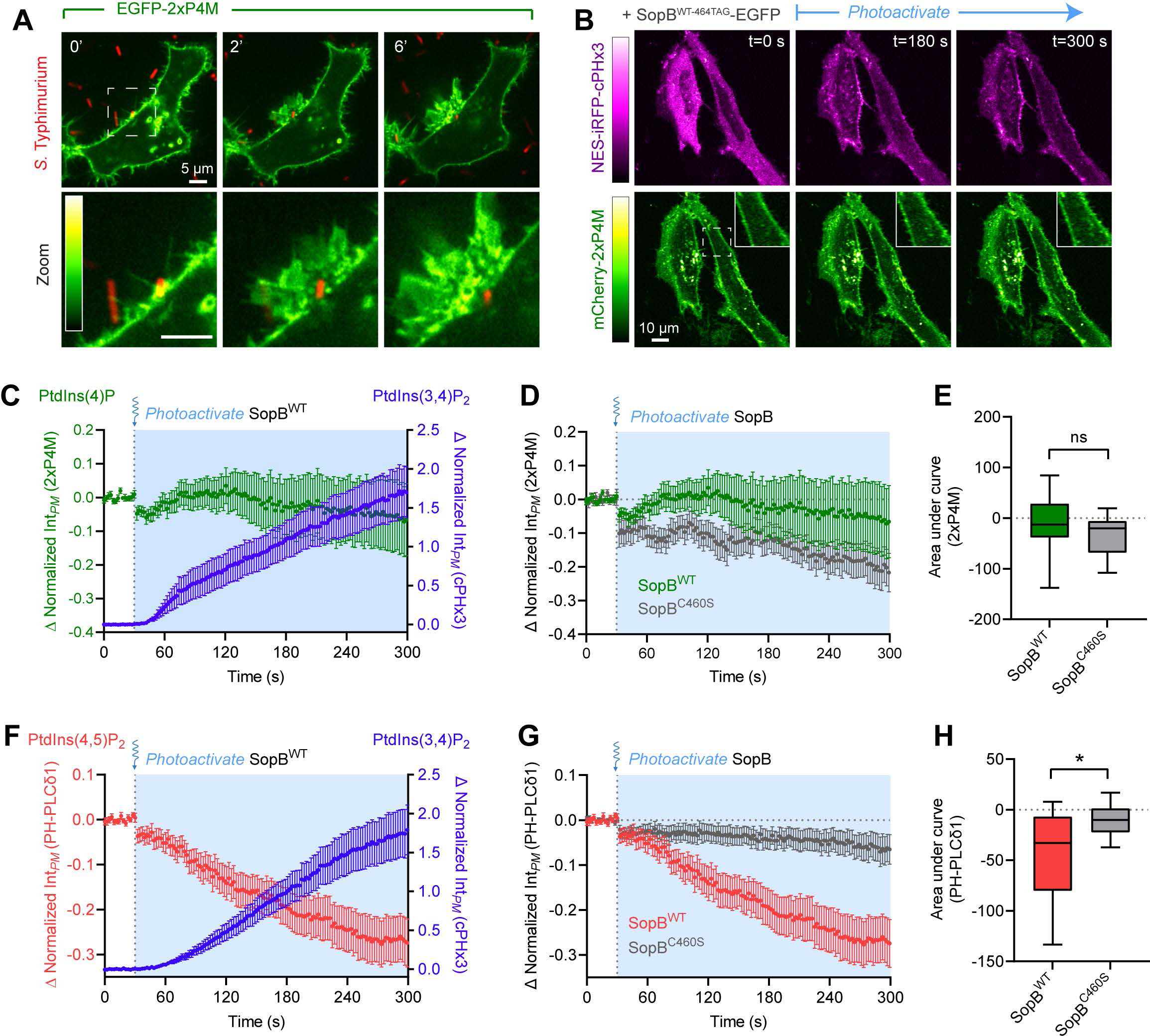
PtdIns(4,5)P_2_ levels correlate inversely with SopB-mediated PtdIns(3,4)P_2_ formation. (A) PtdIns(4)P during bacterial invasion. A representative time-lapse of a HeLa cell expressing 2xP4M (green) during *Salmonella* (red) invasion (0 min indicates time of bacterial contact with the membrane). Panels at bottom show enlarged region denoted by white hashed box. PtdIns(4)P remains abundant within ruffles but is rapidly cleared from the forming SCV. (B) Plasmalemmal PtdIns(4)P is stable during optogenetic activation of SopB. Representative micrographs of NES-iRFP-cPHx3 (magenta) and mCherry-2xP4M (green) pre- and post-photoactivation of wild-type SopB^464TAG^. Illumination of the sample with 405-nm light began at t=30 sec. Insets show enlarged region denoted by white hashed box . (C) Quantification of experiments like (B) plotting baseline-corrected 2xP4M (green) and cPHx3 (blue) intensities during photoactivation of wild-type SopB^464TAG^ from 3 trials (21 cells). Here and throughout this figure, time lapse data are means ± SEM. (D) Comparison of baseline-corrected 2xP4M responses to photoactivation of SopB^WT-464TAG^ (green) and SopB^C460S-464TAG^ (gray). (E) Area under curve calculations of baseline-corrected 2xP4M intensities from (D). Data are box (25-75^th^) and whisker (10-90^th^) percentile plots of >20 individual measurements compiled from 3 trials (SopB^WT-464TAG^, 21 cells; SopB^C460S-464TAG^, 27 cells). Significance assessed by one-tailed unpaired t-test with Welch’s correction (ns, p = 0.064). (F) PtdIns(4,5)P_2_ depletion during PtdIns(3,4)P_2_ synthesis. Quantification of baseline-corrected PH-PLCδ1 (red) and cPHx3 (blue) intensities in the PM before and after photoactivation of wild-type SopB^464TAG^. Data from 3 independent trials (40 cells). (G) Data from (F) comparing the response of PH-PLCδ1 to photoactivation of SopB^WT-464TAG^ (red) and SopB^C460S-464TAG^ (gray). (H) Area under curve calculations of baseline-corrected PH-PLCδ1 intensities from (G). Data from 3 trials are plotted as in (E) (SopB^WT-464TAG^, 40 cells; SopB^C460S-464TAG^, 10 cells). One-tailed Mann-Whitney test of ranks (*, p = 0.0297).

We therefore turned our attention to the other major PPIns species of the PM, namely PtdIns(4,5)P_2_, by expressing a fluorescent conjugate of the PH domain of human PLCδ1 (Stauffer et al., 1998; Várnai and Balla, 1998). Photo-activation of SopB caused a sharp decrease in plasmalemmal PtdIns(4,5)P_2_ that coincided temporally with the appearance of PtdIns(3,4)P_2_ (Figure 6F). Like the accompanying synthesis of PtdIns(3,4)P_2_ synthesis, the decline in PtdIns(4,5)P_2_ was dependent on the integrity of the cysteine residue at the catalytic site of SopB (Figures 6G and 6H). Similar results were obtained with heterologous expression of IpgD (Figures S3B and S3C). These observations suggested that the decline in PtdIns(4,5)P_2_−previously noted to occur at the base of the invasion ruffle (Terebiznik et al., 2002)− might be related and required for the generation of PtdIns(3,4)P_2_.

To directly probe its requirement as a precursor for SopB-mediated PtdIns(3,4)P_2_ biosynthesis, we depleted PtdIns(4,5)P_2_ by recruiting PLCβ3 to the PM using a rapamycin-induced heterodimerization system (Varnai et al., 2006; Won et al., 2006). Phospholipase C enzymes are particularly useful in this context due to their low activity towards PtdIns(3,4,5)P_3_ and PtdIns(3,4)P_2_ (Serunian et al., 1989), while extensively depleting cellular PtdIns(4,5)P_2_ when recruited to the PM by Lyn_(11)_-FRB-HA (Figures 7A and 7B). Critically, the pre-recruitment of FKBP-tagged PLCβ3 to the PM, but not of an FKBP-control vector, led to a virtually complete inhibition of PtdIns(3,4)P_2_ formation in *Salmonella-*induced ruffles, as monitored by cPHx3 (Figures 7C and 7D).

**Fig. 7.**
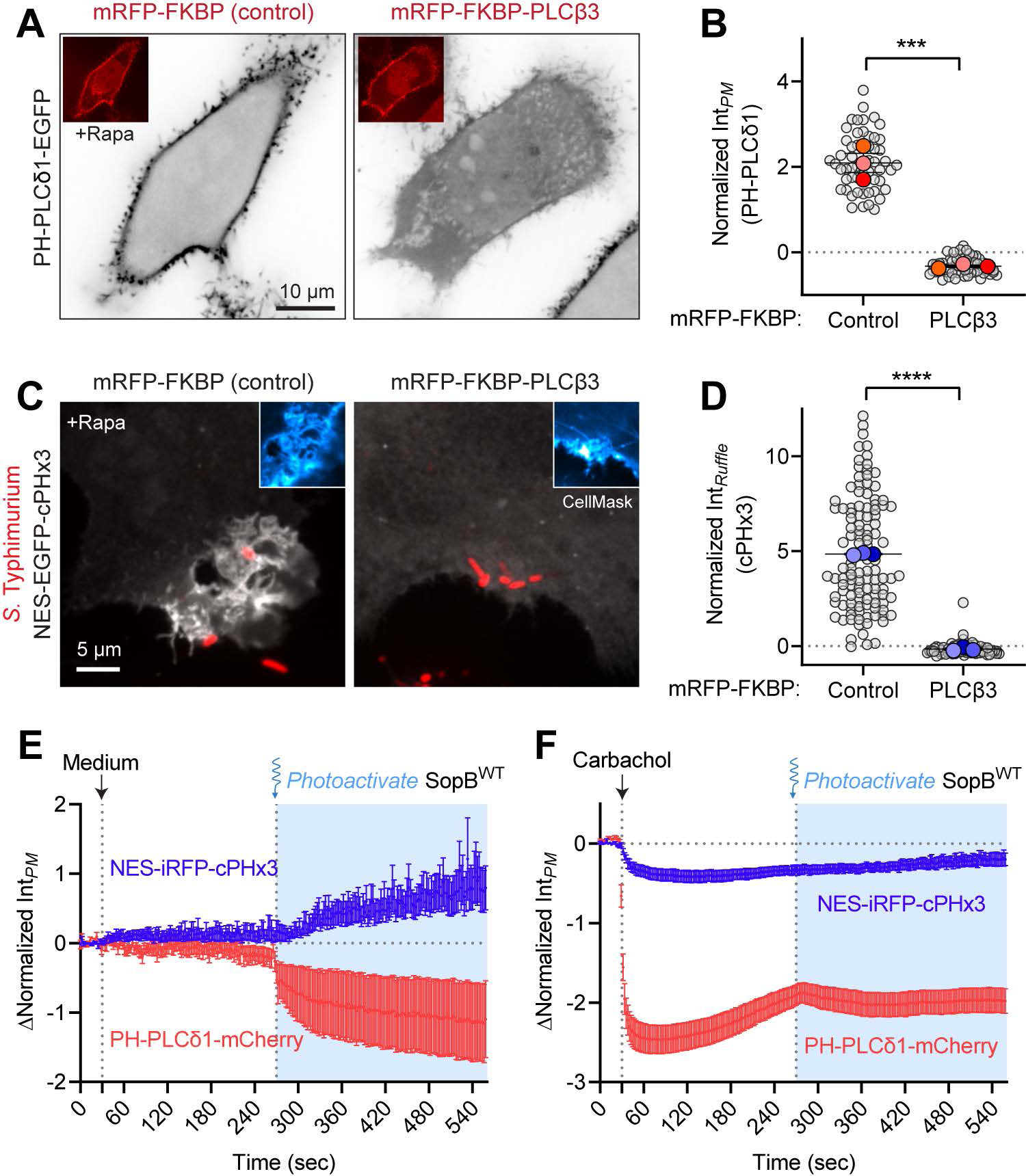
SopB requires a phospholipase C-sensitive inositide to generate PtdIns(3,4)P_2_. (A) Chemically induced recruitment of PLCβ3 to deplete PtdIns(4,5)P_2_ from the PM. HeLa cells expressing mRFP-FKBP (control) or mRFP-FKBP-PLCβ3 together with PM-targeted Lyn_(11)_-FRB-HA and the biosensor PH-PLCδ1-EGFP (inverted gray scale) were treated with 1 µM rapamycin before CellMask staining (not shown) and live imaging. Insets show re-localization of FKBP-conjugates to the PM induced by rapamycin. (B) Plasmalemmal PH-PLCδ1 intensity from experiments like (A) was quantified in 3 trials (control, 55 cells; PLCβ3, 56 cells). Unpaired two-tailed t-test. (C) Inhibition of PtdIns(3,4)P_2_ generated during invasion by pre-recruitment of PLCβ3. Cells transfected as in (A) but expressing the biosensor cPHx3 (gray, extended focus projections) were incubated with 1 µM rapamycin for 2 min before addition of wild-type *Salmonella* (red) for an additional 10 min. The PM was stained (CellMask, inset blue confocal sections of invasion ruffle) before fixation and imaging. (D) The resulting cPHx3 intensities from (C) were quantified in CellMask-labelled invasion ruffles from 3 experiments (control, 115 cells; PLCβ3, 76 cells). Significance assessed by unpaired two-tailed t-test. (E and F) Activation of endogenous PLCβ precludes SopB-mediated PtdIns(3,4)P_2_ synthesis. HeLa cells overexpressing SopB^WT-464TAG^ and muscarinic M3 receptor were subjected to treatment with medium control (E) or 50 µM carbachol (F) at 30 sec to activate endogenous PLCβ, followed by 405-nm light illumination at 270 sec to optogenetically activate SopB. The decrease in PtdIns(4,5)P_2_ and response in PtdIns(3,4)P_2_ were monitored with PH-PLCδ1-mCherry (red) and NES-iRFP-cPHx3 (blue), respectively. Data are means ± SEM of individual cell measurements from 3 experiments. Note the slight decrease in cPHx3 following carbachol treatment likely due to inhibition of class I PI3K signaling constitutively stimulated by the presence of serum.

Because the recruitment of PLCβ3 prior to the addition of *Salmonella* can alter the ruffling induced by the bacteria (insets, Figure 7C) and their ability to invade the host cells, we also evaluated the role of PtdIns(4,5)P_2_ using the photoactivatable SopB. In this instance, we took advantage of endogenous PLCβ enzymes (coupled to G_αq_ subunits) that were activated by addition of carbachol to cells over-expressing the stimulatory M3 muscarinic receptor (Willars et al., 1998). Thus, carbachol-mediated depletion of cellular PtdIns(4,5)P_2_ could be timed with the optogenetic activation of SopB (Figure 7F). Pre-treatment with carbachol strongly depleted PtdIns(4,5)P_2_, as indicated by the loss of plasmalemmal PH-PLCδ1 (coloured red in Figures 7E-F), and blocked subsequent generation of PtdIns(3,4)P_2_ in response to photoactivated SopB. As before, PtdIns(3,4)P_2_ was formed normally in these cells when carbachol pretreatment was omitted (Figures 7E). Together, these results suggest that SopB consumes PtdIns(4,5)P_2_ as it generates PtdIns(3,4)P_2_ in host cells. However, they do not rule out requirement for additional lipids or host factors.

### SopB functions as a phosphotransferase in vitro

We next considered the possibility that SopB could directly generate PtdIns(3,4)P_2_ through the remodelling of pre-existing PPIns, likely including PtdIns(4,5)P_2_. To this end we purified a recombinant construct consisting of residues 33-554 of SopB, that lacks its predicted unstructured regions but retains the putative catalytic site as well as its catalytic activity *in vivo* (Figures S5A and S5B). To facilitate its purification, the catalytically active SopB fragment was fused to the *S. cerevisiae* protein SMT3 (SUMO) and expressed in *E. coli*. The phosphatase activity of this fusion protein was initially validated measuring the release of inorganic phosphate from small unilamellar vesicles containing PtdIns(4,5)P_2_ and PtdSer; no phosphate was released from vesicles containing PtdSer only, confirming the specificity of the recombinant enzyme towards PPIns (Figure 8A), as reported earlier (Marcus et al., 2001).

**Fig. 8.**
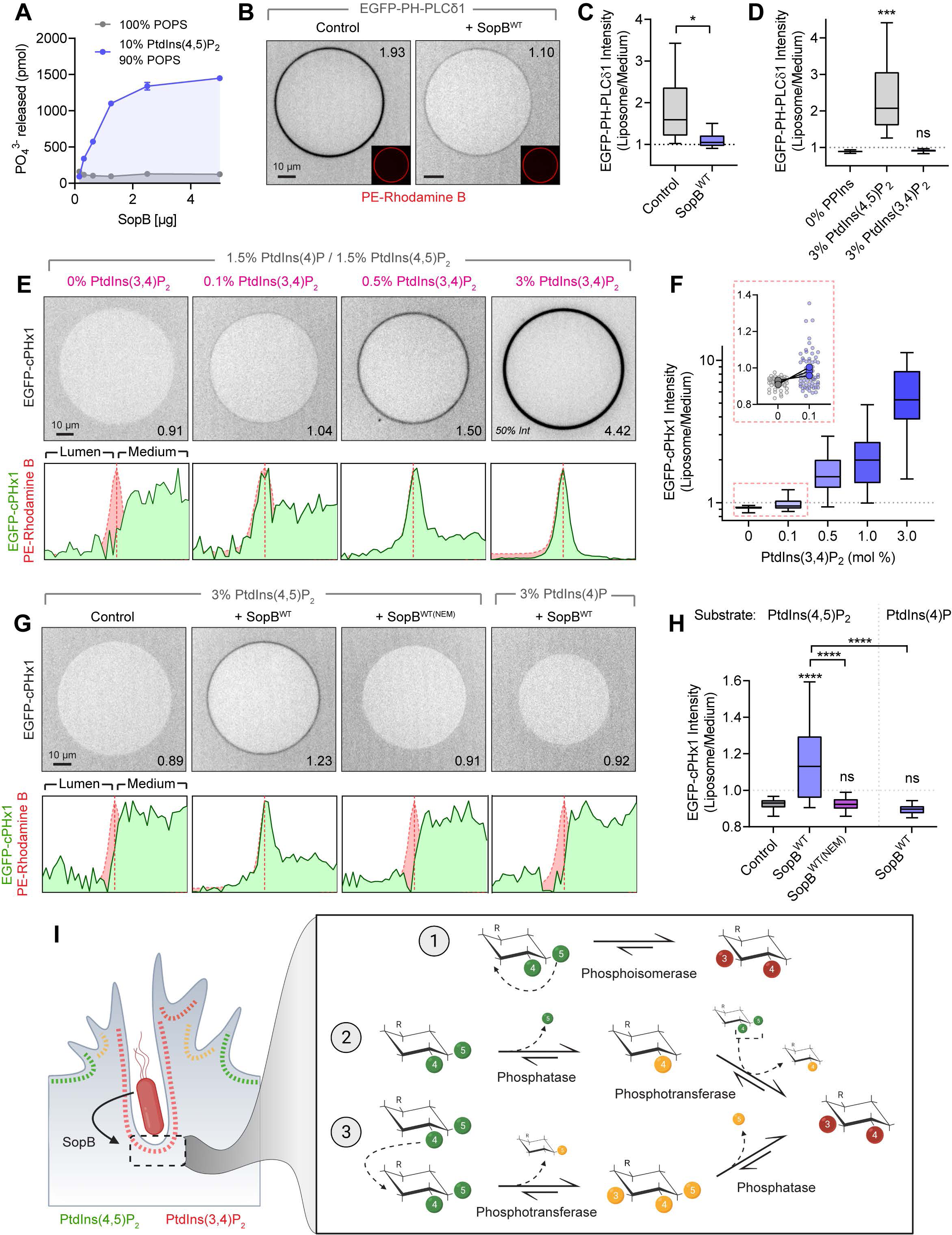
*In vitro* reconstitution reveals SopB phosphotransferase activity. (A) Inorganic phosphate release assessed by malachite green following 30 min treatment of liposomes of the indicated composition with His-SMT3-SopB^33-554^, hereafter SopB. Data are means ± SEM from a representative experiment. (B) Confocal micrographs of GUVs treated for 30 min with SopB or an equal volume of dialysis buffer (control). Liposome composition was PtdCho: PtdSer: PtdIns(4,5)P_2_: PtdEth-Rhodamine B: DSPE-PEG-Biotin (77: 20: 3: 0.1: 0.1 in mol %). Recombinant EGFP-PH-PLCδ1 was added prior to microscopy. See methods for details. (C) Normalized EGFP-PH-PLCδ1 liposome intensity from 4 independent experiments (control, 142 GUVs; SopB, 155 GUVs). Here and elsewhere, graphs are box (25-75^th^ percentile) and whisker (5-95^th^ percentile) plots of individual liposome measurements. Significance determined by two-tailed ratio paired t-test of trial means (*, p=0.0269). (D) Normalized EGFP-PH-PLCδ1 liposome intensity from 3 independent experiments (0% PPIns, 86 GUVs; 3% PtdIns(4,5)P_2_, 84 GUVs; 3% PtdIns(3,4)P_2_, 83 GUVs). One-way ANOVA with Bonferroni’s multiple comparison test of trial means versus control. (E) Representative micrographs of recombinant EGFP-cPHx1 incubated with increasing mol % PtdIns(3,4)P_2_. Background composition of liposomes was PtdCho: PtdIns(4,5)P_2_: PtdIns(4)P: PtdIns(3,4)P_2_: PE-Rhodamine B: DSPE-PEG-Biotin (77-*X*: 1.5: 1.5: *X*: 0.1: 0.1). Corresponding intensity profiles of EGFP (green) and Rhodamine B (red) channels are plotted below. (F) Quantification of liposome-associated EGFP-cPHx1 intensity (part E) from 3 experiments (0%, 53 GUVs; 0.1%, 64 GUVs; 0.5%, 80 GUVs; 1.0%, 68 GUVs; 3.0%, 78 GUVs). Inset graph (red hashed box) shows individual measurements (smaller background circles) and paired trial averages (larger foreground circles) from 0% and 0.1% PtdIns(3,4)P_2_-containing GUVs. (G) GUVs treated for 30 min with dialysis buffer (control), SopB (WT), or SopB (WT-NEM, enzyme pre-treated with a molar excess of NEM), were analyzed by confocal microscopy. Representative EGFP-cPHx1 localization is presented with corresponding normalized values and intensity profiles. GUV compositions (mol %) were PtdCho: PtdSer: PtdIns(4,5)P_2_ or PtdIns(4)P: PtdEth-Rhodamine B: DSPE-PEG-Biotin (77: 20: 3: 0.1: 0.1). (H) Quantification of part (G) from 4 to 5 independent trials per condition (Control, 177 GUVs; SopB^WT^, 185 GUVs; SopB^WT(NEM)^, 147 GUVs; SopB^(WT)^ PtdIns(4)P substrate, 130 GUVs). Significance assessed by one-way ANOVA with Bonferroni’s multiple comparison test of trial averages. (I) Schematic illustration of possible pathways of PtdIns(4,5)P_2_ to PtdIns(3,4)P_2_ conversion by the phosphoinositide phosphotransferase activity of SopB. During invasion (left), PtdIns(3,4)P_2_ accumulates both in PM ruffles and the invaginating regions of the PM prior to fission of the forming vacuole neck. PtdIns(4,5)P_2_ suffices to generate PtdIns(3,4)P_2_ via three possible phosphotransfer-based mechanisms (right): 1) intramolecular transfer (phosphoisomerase), 2) intermolecular transfer preceded by phosphatase activity, and 3) intermolecular transfer followed by phosphatase activity. The latter pathways differ in the predicted intermediate species.

The activity of the recombinant SopB was also validated using giant unilamellar vesicles (GUVs), which are amenable to analysis by confocal microscopy when combined with fluorescent lipid probes. GUVs were immobilized onto streptavidin-coated coverslips by incorporating biotinylated PtdEth (DSPE-PEG-Biotin) and initially located by incorporating 0.1% rhodamine B-conjugated PtdEth, and phosphoinositide changes were monitored as the association or dissociation of soluble regio-isomer-specific EGFP-tagged probes. As anticipated, purified recombinant EGFP-PH-PLCδ1 bound to the outer surface of GUVs containing PtdIns(4,5)P_2_, but not to those containing an equimolar amount of PtdIns(3,4)P_2_, confirming the specificity of the probe (Figure 8D). Importantly, the binding of EGFP-PH-PLCδ1 diminished markedly when GUVs containing PtdIns(4,5)P_2_ were preincubated with SopB (Figures 8B, 8C).

Considering the preceding observations, it was conceivable that SopB was itself converting PtdIns(4,5)P_2_ or a product of its hydrolysis into PtdIns(3,4)P_2_. To address this possibility experimentally, we generated and purified recombinant EGFP-cPHx1 to probe the formation of PtdIns(3,4)P_2_ in a cell-free system. The recombinant EGFP-cPHx1 (containing a single cPH domain) bound to GUVs in a manner proportional to the concentration of PtdIns(3,4)P_2_; no binding was detected in the absence of PtdIns(3,4)P_2_, even when other PPIns species were present (Figures 8E and 8F). The lower limit of detection of PtdIns(3,4)P_2_ with this recombinant sensor was 0.1 mol% (see inset graph, Figure 8F). Using this system we proceeded to test whether SopB was capable of generating PtdIns(3,4)P_2_ in a recombinant system of defined composition. Liposomes containing PtdCho:PtdSer:PtdEth-Rhodamine B:DSPE-PEG-Biotin, (76.8:20:0.1:0.1) plus 3% of the phosphoinositide of interest were generated in phosphate-free buffer and incubated with SopB as in Figure 8B. EGFP-cPHx1 was added to the mixture immediately before imaging to probe for the possible formation of PtdIns(3,4)P_2_. As shown in Figures 8G and 8H, the addition of recombinant SopB to PtdIns(4,5)P_2_-containing liposomes caused the translocation of EGFP-cPHx1 to the liposome surface. This effect was dependent on SopB activity, as it was abolished by pre-treating SopB with N-ethylmaleimide (NEM), which selectively modifies the thiol group of cysteine residues (Gregory, 1955), including Cys460. The ability of SopB to produce PtdIns(3,4)P_2_ in this assay required PtdIns(4,5)P_2_; pre-incubation of liposomes containing an equimolar amount of PtdIns(4)P with SopB did not result in recruitment of EGFP-cPHx1 (Figures 8G and 8H, right panels).

It is noteworthy that the cell-free system we used was devoid of high-energy phosphates (e.g. ATP) and of the divalent cations generally required by kinases. Thus, we concluded that, in addition to functioning as a polyphosphoinositide phosphatase in host cells, SopB possesses phosphotransferase activity, relocating the phosphate groups of PtdIns(4,5)P_2_ to yield PtdIns(3,4)P_2_. Whether this enzyme acts as an intramolecular transferase (i.e., a phosphoisomerase) or an intermolecular transferase (Figure 8I) remains unclear.

## Discussion

Several species of the *Enterobacteriaceae* family diverged from harmless symbionts to parasites. A key evolutionary driver of this feat was the acquisition of virulence factors that facilitate cellular adhesion, bacterial uptake, and manipulation of host signaling (LaRock et al., 2015; Schroeder and Hilbi, 2008). The activation of the host pro-survival kinase Akt by a *Salmonella* effector was described over 20 years ago (Steele-Mortimer et al., 2000). This startling finding spurred the subsequent identification of analogous effector activity secreted by *Shigella* (Marcus et al., 2001; Pendaries et al., 2006). By regulating the survival and proliferation of the infected host cells, Akt plays a key role in the complex host-pathogen inter-species interaction. Its activation counters the pro-apoptotic nature of co-secreted effectors, thereby promoting intracellular growth of the bacteria (Finn et al., 2017; Knodler et al., 2005; Pendaries et al., 2006); Akt activation also alters the inflammatory response (Bruno et al., 2009; Hu et al., 2017; Kum et al., 2011; Zhang et al., 2018) and drives the expansion of the invasion-targeted M cell population (Tahoun et al., 2012). Importantly, Akt activation by the bacteria imprints oncogenic cellular phenotypes during infection-associated carcinoma (Scanu et al., 2015). Akt was known to be activated by PtdIns(3,4,5)P_3_ or PtdIns(3,4)P_2_ (Ebner et al., 2017; Liu et al., 2018), but how accumulation of the responsible 3-PPIns lipid(s) is induced by the enteropathogens had not been defined.

Our data indicated that SopB of *Salmonella* and IpgD of *Shigella* are sufficient to acutely stimulate the synthesis of PtdIns(3,4)P_2_ in host cells. Remarkably, they accomplish this without need to harness any of the previously recognized 3-phosphoinositide biosynthetic pathways such as class I, class II, or class III PI3Ks. Rather, the lipid arises by an effector-catalyzed phosphotransferase reaction. Several lines of evidence indicate that PtdIns(4,5)P_2_ is the sole substrate required for the generation of PtdIns(3,4)P_2_. First, PtdIns(3,4)P_2_ production during infection is restricted to the plasma membrane (Figure 1), the main PtdIns(4,5)P_2_ reservoir in the host cells. Second, the invariable cysteine residue (C460 in SopB, C439 in IpgD) that is essential for the effector-induced disappearance of PtdIns(4,5)P_2_ is also essential for PtdIns(3,4)P_2_ production (Figures 2, 6, and S3). Third, depletion of PtdIns(4,5)P_2_ by heterodimerization-induced recruitment of PLC obliterated the generation of PtdIns(3,4)P_2_ by SopB (Figure 7). Fourth, and most important, the presence of PtdIns(4,5)P_2_ –but not of other phospholipids, including other phosphoinositides– was absolutely required for SopB to generate PtdIns(3,4)P_2_ in a cell-free reconstituted system using recombinant protein (Figure 8).

SopB had earlier been reported to function as a phosphoinositide phosphatase (Anderson Norris et al., 1998; Marcus et al., 2001), an observation we were readily able to confirm by measuring the release of phosphate from PtdIns(4,5)P_2_-containing liposomes (Figure 8A). This activity involves Cys460, which resides within a cationic Cys-X_5_-Arg motif and resembles the sequence of similarly cationic substrate binding clefts found in the phosphate-binding loop of the protein tyrosine phosphatase superfamily (Hsu and Mao, 2015). It is noteworthy that Cys460 is also essential for the generation of PtdIns(3,4)P_2_ from PtdIns(4,5)P_2_; we therefore propose that a common intermediate step underlies the phosphatase and phosphotransferase activity of SopB (and presumably also IpgD). In the case of simple phosphatases, cysteine-mediated nucleophilic attack of a scissile phosphate generates a cysteinyl-phosphate intermediate that is normally hydrolyzed by water to complete the phosphatase cycle (Denu and Dixon, 1998). We envisage three possible catalytic mechanisms to account for the observed phosphotransferase reaction (Figure 8I): Cys460 could act as a nucleophile, attacking one of the phosphates (e.g. D-5) on PtdIns(4,5)P_2_, breaking the bond between the phosphate and inositol ring, and forming a high-energy phospho-cysteine intermediate. Rather than abstracting a proton from water, as would occur during an ordinary phosphatase reaction, proton abstraction would occur from the D-3 position of the inositol ring, promoting its nucleophilicity. The neighbouring Asp465 of SopB could conceivably donate a proton to the leaving group (D-5) of the inositol ring. Thus, the D-3 hydroxyl group nucleophile would be prompted to attack the phospho-cysteine intermediate, generating PtdIns(3,4)P_2_ and regenerating the free cysteine residue. By catalyzing such an intramolecular rearrangement, SopB would operate as a phosphoisomerase (reaction 1, Figure 8I).

Transfer of a phosphate from one inositide to another (i.e. intersubstrate) by an analogous mechanism is also plausible (reactions 2 and 3; Figure 8I) and could operate in combination with the phosphatase activity of SopB. By this putative mechanism, the phospho-cysteine intermediate would attach a second phosphate group onto position D-3 of PtdIns(4)P, generated previously by dephosphorylation of PtdIns(4,5)P_2_. This would require sequential phosphatase and phosphotransferase reactions. Lastly, the reverse sequence can also be contemplated (reaction 3, Figure 8I): the phospho-cysteine intermediate could insert an additional phosphate into PtdIns(4,5)P_2_, generating PtdIns(3,4,5)P_3_. This phosphotransfer reaction would then be followed by dephosphorylation at position D-5, yielding PtdIns(3,4)P_2_. PtdIns(3,4,5)P_3_ could thus be a fleeting intermediate in the biogenesis of PtdIns(3,4)P_2_ by SopB, accounting for the minute amount of PtdIns(3,4,5)P_3_ detected by biochemical measurements during *Salmonella* infection (Mallo et al., 2008). This particular inter-substrate mechanism would also be consistent with the generation of PtdIns(5)P previously identified in host cells (Mason et al., 2007).

The exact residues contributing to the architecture and catalysis in the SopB active site are difficult to discern without precise structural information, which is as yet unavailable; it is notable, however, that deletion of residues 520 to 554 of SopB completely abrogated the phosphotransferase activity (Figure S5C). Cationic residues encoded within this region (K516/525/528/541/553 and R542) could conceivably form an electrostatic counterpart to stabilize the negatively charged inositol head group. Supporting this notion, two of these sites modified in a mutagenesis screen decreased Akt activation in response to SopB (Marcus et al., 2001).

The accumulation of PtdIns(3,4)P_2_ correlated with the phosphorylation of Akt (Figures S1E and S1F). Interestingly, earlier studies identified that Akt2 was the predominant isoform activated by SopB during infection *in vitro* (Cooper et al., 2011) and also *in vivo* (Kum et al., 2011). Consistent with this observation, recent studies suggest that Akt2 is preferentially responsive to PtdIns(3,4)P_2_, while Akt1 and Akt3 are much less sensitive to this inositide and only achieve maximal activation in the presence of PM PtdIns(3,4,5)P_3_ (Braccini et al., 2015; Liu et al., 2018). Thus, the bacteria preferentially target a specific isoform during gastroenteritis.

In summary, this work identifies an unprecedented PtdIns(4,5)P_2_ to PtdIns(3,4)P_2_ conversion mechanism mediated by a phosphotransferase. Through a remarkable instance of convergent evolution, pathogenic organisms have acquired the ability to manipulate host signaling pathways that are central to endocytosis and survival signaling, and can even provoke cancerous cell transformation, as illustrated by the development of gallbladder carcinomas associated with *Salmonella* infection (Scanu et al., 2015).

## Methods

### Plasmids and siRNA

Plasmids utilized in this study are summarized in Table 1. Plasmids were constructed using In-Fusion HD EcoDry Cloning Kits (Takara 121416), NEB HiFi assembly (New England Biolabs E5520S), or traditional restriction enzyme methods and were verified by Sanger sequencing.

**Table 1.**
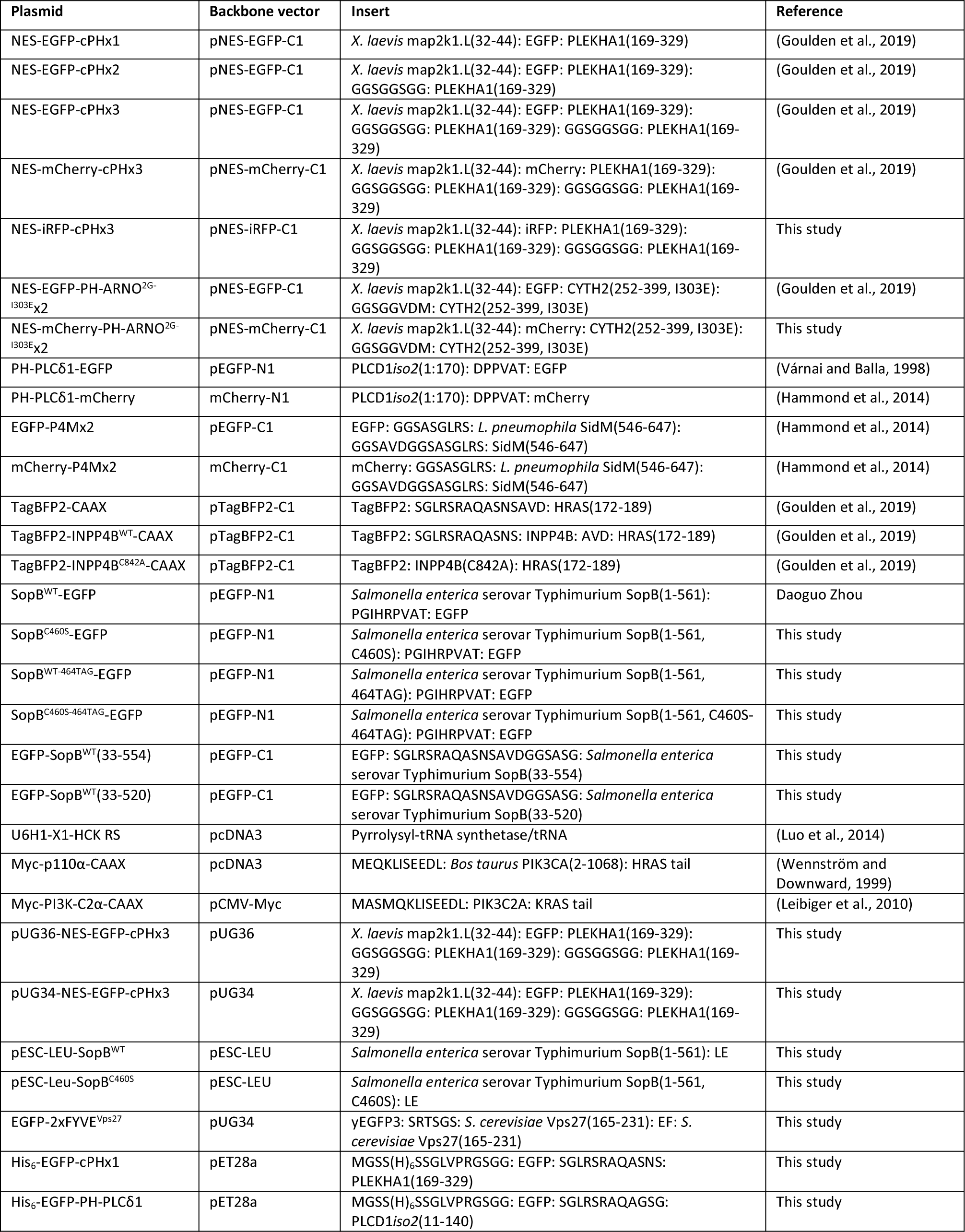
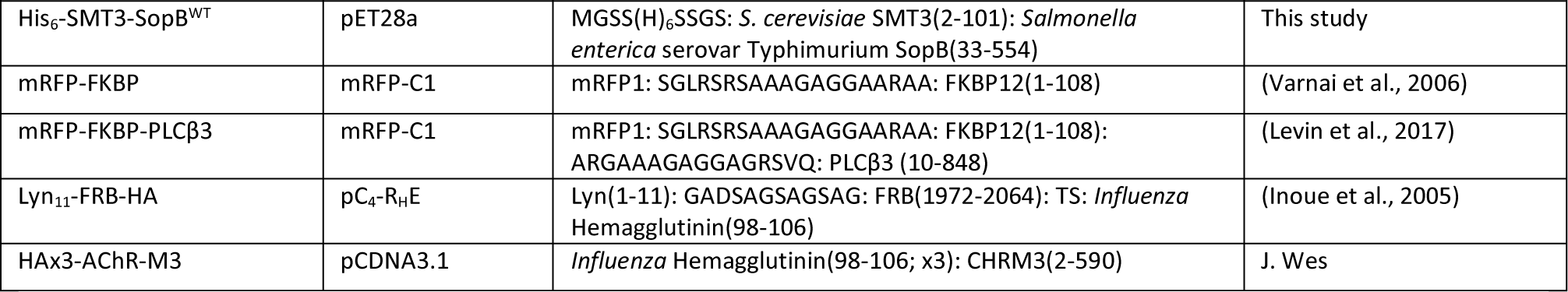
Plasmids used in this study. All genes are of human origin unless otherwise stated.

Mammalian expression vectors are based on the Clontech pEGFP-N1 and pEGFP-C1 vectors with exceptions below. For multichannel imaging, EGFP (from *Aequorea* Victoria; codon-optimized with F64L and S65T mutations) (Cormack et al., 1996), mRFP1 and mCherry (from *Discosoma sp.*) (Campbell et al., 2002; Shaner et al., 2004), the near-infrared fluorescent protein iRFP (from *Rhodopseudomonas palustris*) (Filonov et al., 2011), and the GFP-like protein eq578 variant TagBFP2 (from *Entacmaea quadricolor*) (Subach et al., 2011) were utilized. Site-directed mutagenesis was performed by PCR amplification using targeted pairs of oligonucleotides centrally housing the mutation. The codon corresponding to lysine 464 of wild-type and C460S SopB was substituted with the infrequently used Amber STOP codon (TAG) for photoactivation studies.

Expression of the cPH constructs and the tandem FYVE domains of Vps27 in yeast was achieved by subcloning the open reading frame into pUG34 or pUG36. The cDNA utilized for the generation of pUG34-GFP-2xFYVE^Vps27^ was kindly provided by Scott D. Emr (Zhu et al., 2017).

SopB and the inactive version were cloned into pESC-LEU (Agilent Technologies) for galactose inducible expression.

Recombinant protein expression vectors were constructed from the pET28a series (Novagen) encoding an N-terminal hexa-histidine tag and thrombin cleavage site.

Custom basic RNA oligos (Thermo Fisher 10620310) targeting *PIK3C2A* were 5’-GGAUCUUUUUAAACCUAUU-3’ (sequence 1) and 5’-GCACAAACCCAGGCUAUUU-3’ (sequence 2) described previously (Posor et al., 2013). 5’-phosphorylated oligonucleotides were suspended in water to 20 μM. An equimolar amount of ON-TARGETplus Non-targeting Control Pool (Dharmacon D-001810-10-20) was utilized as control.

### Reagents, Antibodies, and Lipids

Wortmannin (Sigma 681675) was dissolved in DMSO to 5 mM. PI-103 (Sigma 528100) was dissolved in DMSO to 5 mM. GDC-0941 (Millipore Sigma 509226) was dissolved in DMSO to 10 mM. Carbachol (Fisher Scientific AC10824-0050) was dissolved in water to 50 mM. CellMask Deep Red (ThermoFisher C10046) was dissolved in DMSO to 5 mg/mL. Carbenicillin Disodium (Wisent Bio Products 400-112-IG) was dissolved in water to 100 mg/mL. Kanamycin (BioBasic KB0286) was dissolved in water to 50 mg/mL. Gentamycin (Wisent Bio Products 450-135-XL) was dissolved in water to 50mg/mL. Streptomycin (Sigma S9137) was dissolved in water to 50 mg/mL. Rapamycin from *Streptomyces hygroscopicus* (Sigma R0395) was dissolved in DMSO to 5 mM. Streptavidin from *Streptomyces avidinii* (Sigma S4762) was resuspended in PBS to 1 mg/mL. Dithiothreitol (BioBasic Inc. DB0058) was resuspended in water to 1 M. Concanavalin A, Alexa Fluor 647^TM^ conjugate (Invitrogen C21421) was dissolved in 0.1 M Sodium Bicarbonate pH 8.3 to 5 mg/mL. Hydroxycoumarin lysine (Luo et al., 2014) was dissolved in DMSO at 100 mM. All were aliquoted and stored at –20 °C.

PLL(20)-g[3.5]-PEG(5) and PLL(20)-g[3.5]-PEG(2)/PEG(3.4)-biotin(20%) (Susos Surface Technology) were suspended in 150 mM NaCl, 20 mM Tris pH 7.4 to 1 mg/mL and were stored at 4 °C. Ultra-low gelling temperature agarose (Type IX-A; Sigma Aldrich A2576) was suspended by heating in molecular grade water to 1% (w/v) and then stored at 4 °C. Malachite green solution (Echelon Biosciences K-1501) was stored at 4 °C.

The following primary antibodies were utilized during immunofluorescence and/or immunoblotting: Mouse mAb anti-Akt pan (Cell Signalling 40D4, 2920; RRID:AB_1147620); Rabbit mAb anti-Akt pS473 (Cell Signalling D9E, 4060); Rabbit pAb anti-Akt pT308 (Cell Signalling 9275, RRID:AB_329828); Mouse mAb anti-GAPDH (EMD Millipore 6C5, MAB374; RRID: AB_2107445); Rabbit pAb anti-PI3K-C2α (ProteinTech 22028-1-AP; RRID:AB_11183760); Mouse mAB anti-c-Myc (DSHB 9E10, RRID:AB_2266850); Mouse mAb anti-vinculin (EMD Millipore VIIF9 (7F9), MAB3574; RRID: RRID:AB_2304338); Rabbit Ab anti-*Salmonella* O Antiserum Group B Factors 1, 4, 5, 12 (Fisher Scientific DF2948-47-6; RRID:AB_2884995); Mouse mAb anti-Cdc42 (BD Biosciences 610928; RRID:AB_398243); and Mouse mAb anti-α-tubulin (Sigma T5168; RRID:AB_477579). HRP-conjugated secondary antibodies raised in Donkey and all fluorescently conjugated secondary antibodies (DyLight 405, Alexa Fluor 488^TM^, Alexa Fluor 647^TM^) were from Jackson ImmunoResearch Laboratories Inc. Infrared secondary antibodies IRDye 800CW and IRDye 680RD used for immunoblotting were from Li-Cor.

The following lipids were utilized *in vitro*: PtdIns(4,5)P_2_ diC16 (Echelon Biosciences P-4516-2); PtdIns(4)P diC16 (Echelon Biosciences P-4016-2); PtdIns(3,4)P_2_ (Echelon Biosciences P-3416-2); PtdIns(3,4,5)P_3_ (Echelon Biosciences P-3916-2); 16:0-18:1 PC (POPC, Avanti Polar Lipids Inc. 850457C); 16:0-18:1 PS (POPS, Avanti Polar Lipids Inc. 840034; 18:1 Liss Rhod PE (Avanti Polar Lipids Inc. 810150); and DSPE-PEG(2000) Biotin (Avanti Polar Lipids Inc. 880129C). PPIns were solubilized in 1 mL of CHCl_3_:MeOH:1M HCl (2:1:0.01 ratio), vortexed, and dried under N_2_ stream before three resuspension and drying cycles in CHCl_3_. Finally, PPIns were resuspended in CHCl_3_ to a final concentration of 0.5 mg/mL. All lipids were stored at –20 °C.

### Cell culture and transfection

HeLa (wild-type) cells obtained from the American Type Culture Collection (ATCC) were maintained in DMEM with 1.5 g/L sodium bicarbonate & sodium pyruvate (Wisent Bio Products 319-007-CL) supplemented with 10% heat-inactivated fetal bovine serum (Gibco 12483-020).

Cells were incubated at 37^°^C in a humidity-controlled atmosphere at 5% CO_2_ and were passaged two to three times per week by detachment with 0.25% Trypsin-EDTA (Wisent Bio Products 325-043-EL) and dilution 1 in 5 to 1 in 10. HeLa cells were confirmed to be mycoplasma-negative before freeze-down and were utilized between passage 5 and 25 from receipt. For photoactivation experiments, HeLa were cultured in low glucose DMEM (Life Technologies 10567022) containing 10% heat-inactivated fetal bovine serum (Life Technologies 10438-034), penicillin (100 units/mL), streptomycin (100 μg/mL; Life Technologies 15140122) and chemically-defined lipid supplement (1:1000; Life Technologies 11905031).

Cells were enumerated using the Z2 Coulter particle count and size analyzer (Beckman Coulter) and 8.5x10^4^ cells were seeded two days prior to bacterial invasion onto uncoated 18 mm Number 1.5 glass coverslips (Fisher Scientific 12545100) in 12-well plates. Per two wells of cells seeded 6-24 h prior to transfection, 1.5 μg total DNA was combined with 4.5 μL FuGENE 6 (Promega E2691) pre-complexed in 100 μL serum-free DMEM. Cells were maintained in serum containing medium prior to infections unless otherwise stated in Figure Legends.

Photoactivation experiments were performed by transfecting HeLa cells with plasmids encoding SopB^WT-464TAG^-EGFP (or mutant SopB^C460S-464TAG^), the plasmid encoding the engineered pyrrolysyl-tRNA synthetase/tRNA pair that incorporates HCK in response to an amber stop codon, and various biosensors. For unnatural amino acid incorporation, medium was supplemented with 250 µM of hydroxycoumarin lysine (HCK) in parallel or 4 hours after transfection. In response to the UAG codon encoded in *sopB*, HCK is transacylated onto the tRNA and ribosomal incorporation of HCK in the growing polypeptide occurs in lieu of lysine 464 of SopB.

For RNA interference, HeLa cells were seeded directly in 12-well plates at a density of 1.1x10^5^ cells/well and were transfected on day 2 with 100 nM siRNA in Opti-MEM (Gibco 31985-070) using Lipofectamine RNAiMAX (Thermo Fisher 13778030) according to the manufacturer’s instructions. Media was changed on day 3 prior to a second round of siRNA transfection. Cells were lifted on day 4, counted, and seeded in parallel for immunoblotting (1.6x10^5^ cells/well) and microscopy experiments (1.3x10^5^ cells/coverslip). Plasmids encoding biosensors were transfected 6-8 h after re-plating using FuGENE 6 as described above. In parallel on day 5, lysates were collected for immunoblotting from the commonly derived population of cells utilized for bacterial invasion and microscopy.

Imaging commenced 18-24 h post-transfection except for transient transfection of SopB-EGFP (Wild-type, C460S) where 8-12 h yielded sufficient expression, and transient expression of TagBFP2-INPP4B-CAAX (Wild-type, C842A) which we found empirically to yield improved expression 40-48 h post-transfection.

### S. Typhimurium strains, culture, and invasion

*S.* Typhimurium strains used in the current study are derivatives of wild type *Salmonella* enterica subspecies I serovar Typhimurium strain SL1344 (Hoiseth and Stocker, 1981). Isogenic Δ*sopB* SL1344 generated by allelic exchange has been previously described (Steele-Mortimer et al., 2000). Corresponding mRFP1 (Campbell et al., 2002)- and BFP (Heim and Tsien, 1996)-expressing strains were generated by electroporating the low copy plasmids (pFPV25.1 encoding BFP and pBR322 encoding mRFP1) under the control of the *rpsM* promoter (Valdivia et al., 1996). Epithelial cells were exposed to late-log phase *Salmonellla* by a method optimized for bacterial invasion (Steele-Mortimer et al., 1999). Wild-type and isogenic mutants (3-5 colonies) were grown with antibiotic selection overnight at 37 °C. Stationary phase cultures were diluted 1:33 into fresh LB without antibiotics for 3.5 h at 37 °C when inocula were prepared by centrifugation at 10,000 x g for 2 min, and resuspension in an equivalent volume of D-PBS.

To monitor biosensors (statically) during invasion, cells were infected in 1 mL D-PBS by late-log phase *Salmonella* (mRFP-WT, Δ*sopB*; 1:50 dilution) for 10 min at 37 °C, 5% CO_2_. Coverslips were submerged for 5 min in pre-chilled D-PBS containing CellMask Deep Red (1 μg/mL), followed by two gentle washes with chilled D-PBS, and fixation in 2.0-2.5% PFA for 20 min.

Cells were imaged immediately following three PBS washes. For inhibitor experiments, cells were pre-treated in pre-warmed D-PBS for 20 min with concentrations indicated within individual Figure Legends. Inhibitors were maintained in the medium throughout bacterial invasion. During timecourse studies, extracellular bacteria were removed by extensive washing in D-PBS and by the inclusion of 100 μg/mL gentamycin in media 0.5 h post-infection.

Gentamycin was reduced to 10 μg/mL at 2 h post-infection for the remainder of the experiment.

### Yeast strains, growth media, and genetic methods

SEY6210 wild-type *S. cerevisiae* (*MATα leu2-3,112 ura3-52 his3-Δ200 trpl-Δ901 lys2-801 suc2-Δ9 GAL*) (Robinson et al., 1988) and SEY6210 *vps34Δ1::TRP1* (PHY102) (Herman and Emr, 1990) transformed with a low-copy plasmid (*CEN* URA3) encoding a temperature-sensitive mutant of the *vps34* allele (Stack et al., 1995) have been previously described. Strains were transformed with plasmids indicated in Table 1 by the lithium-acetate method (Gietz and Woods, 2002) and maintained on appropriate minimal medium containing 1.7 g/L yeast nitrogen base without amino acids, 5 g/L ammonium sulphate, amino acids, 2% (w/v) D-Glucose or 2% (w/v) D-Galactose, and 2% (w/v) bacto-agar. Yeast strains grown overnight in appropriate media at 28-30 °C were re-inoculated into fresh glucose-containing medium at 0.25 OD_600_ units and allowed to grow for 3 h. The logarithmically growing cells were then transferred into galactose-containing medium for 2 h to induce SopB before imaging. Vps34 was inactivated in the *vps34^ts^* strain by transferring the cells to 38 °C (non-permissive temperature) for 90 min prior to inducing SopB with galactose at the same temperature for 60-90 min. Cells were collected by centrifugation and were resuspended in medium containing 0.1% Trypan Blue prior to mounting on 2% agarose (w/v) pads overlaid with Number 1.5 1.8-cm glass coverslips. Where indicated, Concanavalin A (5 μg/mL) was added for 10 min prior to centrifugation to demarcate the cell wall.

### Immunostaining

Paraformaldehyde (2% w/v)-fixed cells were permeabilized in 0.1 % (v/v) Triton X-100 in PBS for 5 min, blocked in 2% (w/v) BSA in PBS for 20 min, and coverslips were overlaid consecutively for 1 h with primary antibodies and secondary antibodies in 1% BSA, separated by PBS washes and a wash in 2% BSA.

### Recombinant protein production

Recombinant His-tagged proteins indicated in Table 1 were expressed and purified from BL21 (DE3) *E. coli* (NEB C2527) using isopropyl β-D-1-thiogalactopyranoside (IPTG)-inducible expression systems and amino-terminal hexa histidine tags. Several colonies were inoculated into Terrific Broth containing 4 mL/L glycerol (Multicell Cat No. 800-067-LG: total volume 5 mL) with 50 ug/mL Kanamycin selection and 0.8 % (w/v) D-Glucose. Starter cultures were incubated shaking overnight at 37 °C before 1:50 dilution into pre-warmed, fresh Terrific Broth (300 mL in 1 L Erlenmeyer flask) with antibiotic selection. Cultures were grown at 37 °C to 0.6 to 0.8 OD_600_ units before equilibrating at room temperature (30 min) and inducing expression by the addition of 0.05-0.1 mM IPTG. Cultures were induced for 20 h at 23 °C (210 rpm) before harvesting by centrifugation (5,000 x g for 15 min, 4 °C) and freezing at -20 °C prior to downstream purification.

For the EGFP-tagged TAPP1 cPH domain, half of the resultant pellet was resuspended in 20 mL of 150 mM NaCl, 20 mM Tris, pH 7.4 (at 10 °C) with EDTA-free protease inhibitors (Thermo Fisher A32955) and DNase I (Sigma-Aldrich DN25). Entire pellets of EGFP-tagged PLCδ1 PH domain and SMT3-SopB^33-554^ were resuspended in 25 mL as above but contained 300 mM NaCl, 40 mM Tris, 15 mM Imidazole, 5% (v/v) glycerol, pH 7.4 (at 10 °C), with 5 mM β-Mercaptoethanol, protease inhibitors, and DNase. Resuspended bacteria were disrupted three times using a French press apparatus (SLM AMINCO Spectronic Instruments) before centrifugation of the resultant slurry (15,000 x g for 20 min, 4 °C). Clarified supernatants were incubated with TALON® Metal Affinity Resin (Takara 635502) and rotated at 4 °C for 30 min (cPH domain) to overnight (PLCδ1 domain and SopB). Resin was washed twice with 10-15 mM Imidazole in the corresponding lysis buffer as indicated by the manufacturer, before loading into a TALON® 2 ml Gravity Column (Takara 635606). Columns were washed one additional time before elution in a single step (150 mM Imidazole, TAPP1 cPH domain) or a stepwise gradient (15, 30, 40, 50, 60, 180, 180, 240 mM Imidazole; PLCδ1 PH domain and SopB).

Elution fractions were analysed on 12% polyacrylamide gels and fractions containing products of the predicted molecular weights were pooled (His_6_-EGFP-cPHx1, 48.39 kDa; His_6_-EGFP-PH-PLCδ1, 45.41 kDa; His_6_-SMT3-SopB^33-554^, 69.95 kDa). The identity of His_6_-SMT3-SopB^33-554^ was also confirmed by its sensitivity to Ulp1-mediated proteolysis (product reduced to ∼57 kDa), though after confirming SopB activity towards inositol lipids by phosphate release, the uncleaved His_6_-SMT3-SopB^33-554^ recombinant protein was utilized. Peak His_6_-EGFP-cPHx1 fractions were dialyzed several hundred-fold into imidazole-free buffer (150 mM NaCl, 20 mM Tris, pH 7.4) while concentrating the product to 2.5 mg/mL with serial spins (5,000 x g for 20 min, 4 °C) using Amicon® Ultra-15 Centrifugal Filter Unit, 30 kDA MWCO (Millipore UFC903008). Peak His_6_-EGFP-PH-PLCδ1 fractions were dialyzed (ThermoFisher 66380; Slide-A-Lyzer™ Dialysis Cassettes, 10K MWCO) into imidazole-free buffer (150 mM NaCl, 20 mM Tris, 5% (v/v) glycerol,

0.5 mM β-Mercaptoethanol, pH 7.4 at 10 °C) to a final concentration of 7 mg/mL. Peak His_6_-SMT3-SopB^33-554^ fractions were diluted with additional imidazole-free buffer (300 mM NaCl, 40 mM Tris, 5% (v/v) glycerol, pH 7.4 at 10 °C, 5 mM β-Mercaptoethanol) before dialysis to a final concentration of 0.4 mg/mL. Final products were aliquoted before snap freezing in N_2(l)_ and long-term storage at -80 °C.

### Unilamellar Vesicles and Phosphate Release Assays

Phosphatase activity was assayed using malachite green (Maehama et al., 2000). Lipid mixtures (100% POPS or 90% POPS/10% PtdIns(4,5)P_2_) were suspended by probe sonication on ice to 1 nmol lipid/μL assay buffer, and 25 μL of the suspension was utilized per 96-well. This provided a 1250 pmol theoretical maximum assuming hydrolysis of a single position of the inositol ring and equal bilayer distribution. Reactions were performed for 30 min at 37 °C in the presence of 10 mM DTT, before the addition of malachite green solution (100 μL) and measurements of absorbance at 620-nm. Data were plotted after calibrating absorbance measurements with KH_2_PO_4(aq)_ in 100% POPS.

Giant unilamellar vesicles (GUVs) were generated on a film of agarose as previously described (Horger et al., 2009). Glass slides were overlaid with 1% (w/v) ultra-low gelling temperature agarose (Type IX-A) and dried by heating (40 °C) until translucent. Various liposome compositions were prepared by adding chloroform-suspended lipid stocks to glass vials, drying under N_2(g)_, and resuspending in a small volume of chloroform. Generally, 4/3 (∼1.3) μmol of total lipid were used per preparation (compositions are indicated within individual Figure Legends). Dried lipids were re-suspended in 35 μL chloroform before spreading evenly over the dried agarose. Unused lipid mixes (large batch) were alternatively dried and stored long-term at -20 °C. Chloroform was thoroughly evaporated from the agarose film under N_2(g)_ stream before submerging the slide in a Petri dish containing assay buffer (200 mM NaCl, 20 mM Tris, pH 7.4 at room temperature) for 3 h to overnight. GUVs were harvested by pipetting assay buffer over the slide several times. This solution was used directly or centrifuged (4,000 x g, 5 min) to concentrate large vesicles. Preparations were used within 48 h mainly, but long-term storage (< 7 days) was at 4 °C.

To stabilize GUVs during fluorescence microscopy, acetone-washed 18 mm Number 1.5 glass coverslips were passivated (30 min) with a 3:1 mixture of PLL(20)-g[3.5]-PEG(5) to PLL(20)-g[3.5]-PEG(2)/PEG(3.4)-biotin(20%) diluted in assay buffer to 100 μg/mL. Following washes, the passivated glass was overlaid (30 min) with 25 μg/mL streptavidin in assay buffer which coupled to the biotin-containing group. Thus, following washes, GUVs containing 0.1% DSPE-PEG(2000)-Biotin were stabilized on the glass surface (generally after ∼3-5 min) via the sequential biotin-streptavidin-biotin interactions. For enzymatic reactions, 6 μg of SopB (428.9 nM) was pre-incubated for 30 min at 37 °C in assay buffer containing liposomes (∼666 μM, yields vary) with freshly thawed DTT (5 mM). Fluorescent biosensors (1 μM) were added post-hoc before adding the liposome reaction to prepared coverslips for confocal microscopy.

### Fluorescence Microscopy and Photoactivation

Spinning disk confocal microscopy was performed with a Quorum WaveFX spinning disk system (Quorum Technologies Inc.) which consists of an Axiovert 200M microscope (Carl Zeiss), equipped with a 63x/1.4 NA oil objective and 25x/0.8 NA multi-immersion objective multiplied by a 1.53x magnifying lens. Scanning is performed by the CSU10 confocal multi-beam scanner (Yokagawa). Fluorophores were excited consecutively utilizing a four-line laser module (405, 491, 561, 640 nm; Spectral Applied Research) and filtered by a corresponding emission filter wheel (447/40, 515/40, 515 LP, 594/40, 624/40, 670/40). Signal was detected by a back-thinned EM-CCD camera (Hamamatsu ImageEM C9100-13). This system is driven by a motorized XY stage (Applied Scientific Instrumentation) and Piezo Focus Drive (Ludl Instruments). Acquisition settings and capture are controlled by Volocity software v6.2.1 (PerkinElmer).

For live cell imaging, coverslips were mounted within a Chamlide^TM^ magnetic chamber (Live Cell Instrument Inc.) overlaid with pre-warmed phenol red-free Tyrode’s buffer (140 mM NaCl, 5 mM KCl, 2 mM CaCl_2_, 1 mM MgCl_2_, 10 mM D-Glucose, 10 mM HEPES pH 7.4) and maintained at 37 °C using an environmental chamber (Live Cell Instruments Inc.). Late-log phase *Salmonella* were diluted into a small volume of Tyrode’s buffer and were added directly to the imaging chamber after acquisition of uninfected fields. Generally, up to 10 fields were acquired consecutively with 0.3 μm Z-steps to capture invasion ruffles. Pharmacological inhibitors were diluted to 10x concentration in 50 μL Tyrode’s buffer before addition to the imaging chamber.

High-resolution Airyscan microscopy was performed on the LSM880 Airyscan system (Carl Zeiss) which is equipped with a 63x/1.4 NA oil objective and three detectors: two PMT and one 32-channel spectral Airyscan PMT. Laser lines utilized were 488 nm (Argon) 552 nm, and 642 nm under Airyscan mode. Acquisitions are driven by motorized XY stage and Z-Piezo Drive (Carl Zeiss), and settings and capture controlled by Zen Black software (Carl Zeiss).

Optogenetic experiments were performed by confocal microscopy on a Nikon TiE inverted stand with an A1R resonant scan head and fiber-coupled four-line excitation (Ex) LU-NV laser combiner equipped with 405-, 488-, 561-, and 640-nm lines. 8 or 16 frame averages were used to improve signal to noise. A 100× 1.45 NA plan-apochromatic oil immersion objective was used. To prevent crosstalk between channels, blue (emission 425–475 nm) and yellow/orange (emission 570–620 nm) fluorescence were acquired on a separate excitation scan to the green (emission 500–550 nm) and far red (emission 663–737 nm) channels.

Optogenetic activation of SopB was performed after acquiring ∼30 s of data with 405 nm illumination set to zero power, at which time transmission was increased to 20% of the maximum available power from the LU-NV unit. Cells were imaged in 1.6 ml FluoroBrite DMEM (Life Technologies A1896702) supplemented with 25 mM Hepes (pH 7.4) 1:1,000 chemically defined lipid supplement with 10% heat-inactivated fetal bovine serum.

### Image Analysis

Images in .nd2 (Nikon) or .mvd2 (Volocity) were exported and analyzed in the open-source image analysis platform Fiji (Schindelin et al., 2012). For quantitative analysis of biosensor intensity in the bulk PM or within invasion ruffles, a recruitment index was produced by normalizing the intensity of the fluorescent protein in the membrane to that of the cytosol (Teruel and Meyer, 2000) after correcting for background intensity of the detector. Intensity data are graphically presented as the Int*_PM_*/Int*_Cytosol_* for photoactivation data and reported as the Int*_PM_* -Int*_Cytosol_*/Int*_Cytosol_* for all other cell culture experiments. During photoactivation assays, normalized data were computed by subtracting the average ratio of frames before photoactivation from the ratio along the time lapse. As described below, the PM was defined by the signal of co-expressed fluorescent FRB-conjugates following their rapamycin-mediated recruitment, by CellMask (an amphipathic molecule containing a lipophilic moiety which inserts into the exofacial leaflet of the PM), or by manually defining a small region of interest (ROI) of the PM based on lipid biosensors enriched at the PM prior to photoactivation time-lapses (PH-PLCδ1, 2xP4M, cPHx3).

For estimation of fluorescence intensity in the bulk PM, a binary mask was generated by à trous wavelet decomposition (Olivo-Marin, 2002; Smal et al., 2010) using a custom-written macro in Fiji similar to a previously implemented method (Hammond et al., 2014). To generate this binary mask, a threshold was applied to the wavelet product that included pixel intensities several fold-times the standard deviation of the wavelet product (factor was empirically determined for each data set but ranged from 0.8•StdDev to 1.5•StdDev). Applying this method to fluorescent FRB-conjugates or CellMask-stained cells produced a consistent binary mask that was utilized to quantify the fluorescence intensity of a given biosensor in the PM compartment. Measurements were reported per cell by manually producing ROIs that encompassed single transfected cells, non-nuclear cytoplasm, and background, and restricting measurements to each ROI.

To capture biosensor recruitment to bacterial invasion ruffles, individual circular ROIs (pixel*_radius_* = 100 or ∼13.2 μm) encompassing single invasion sites at the PM were manually annotated from single confocal micrograph sections. As above, fluorescence intensity of biosensors was quantified within a binary mask generated from parallel acquisition of CellMask. Binary masks of ruffles were generated by first smoothing CellMask images with a Gaussian filter. After converting to an 8-bit gray scale, a local threshold was then applied to images using Bernsen’s method (Bernsen, 1986) which is an adaptive binarization method computed for a sliding circular window (pixel*_radius_* = 3). Biosensor intensity was then analyzed within the circular ROI of the invasion site and normalized to the cytosolic intensity.

For quantitation of fluorogenic probe recruitment to liposomes, an annulus of the liposome surface was generated and compared to an expanded annulus of the surrounding medium using a custom-written macro. Otsu’s method was applied to micrographs (PE-Rhodamine channel) to threshold pixel intensities of individually circled liposomes. This was converted to a binary mask that was filled and then iteratively eroded. The resulting binary shape (conforming to the shape of the liposome) was outlined and dilated which created a 4-pixel wide annulus that overlapped tightly with the liposome surface. Finally, a corresponding ‘medium’ measurement was generated by expanding this annulus 10 pixels (dilations) beyond the liposome surface. Data are presented as the resulting Liposome/Medium intensity ratio per individual liposome.

### Statistical analysis and image presentation

Data were imported into GraphPad Prism 9 software for statistical analysis and the presentation of data. Superplots (Lord et al., 2020) were generated to communicate cellular and invasion-level variability (presented in the background as gray transparent data points) and trial-to-trial variability (overlayed trial averages in the foreground as solid colours). The number of cells analyzed, number of independent trials (performed on separate days), statistical tests, and statistical results are indicated within individual Figure Legends. Cell culture experiments were repeated on separate passages of cells. Except for photoactivation assays, trial averages were subjected to statistical analyses. Shapiro-Wilk normality tests applied to the trial means (N ≥ 3) were consistent with a normal distribution so parametric tests were applied (unpaired two-tailed t-test for 2 groups, and unpaired ordinary one-way ANOVA with Bonferroni’s multiple comparisons test for ≥ 3 groups).

To compare photoactivation datasets, baseline-corrected data were sorted into individual groups of curves each comprising measurements of a normalized ROI across an entire time lapse. The area under the curve (AUC) was calculated for each time lapse, and the resulting net AUCs were sorted by condition to compare groups statistically. Shapiro-Wilk normality tests were applied to each grouping, and parametric or non-parametric t-tests were applied accordingly as described within individual Figure Legends.

Representative images were chosen based on being typical of a phenotype, possessing good signal-to-noise ratio, and where applicable, being quantitatively representative of the dataset (close to the trial averages or median). For direct comparison during *in vitro* studies, quantitative fluorescence values are inset in the representative images. Processing of representative images such as merging and cropping for insets was performed in FiJi. Where necessary for visualization, linear adjustments were made to brightness and contrast across the entire image or alternative lookup tables were applied. Exported RGB format TIFF images were sized once in Adobe Photoshop to the final publication format prior to assembling in Adobe Illustrator.

## Acknowledgements

We thank Dr. Scott Emr (Department of Molecular Biology and Genetics, Cornell University) for kindly sharing plasmids for this study. We thank Dr. Kimberley Lau and Paul Paroutis (The Imaging Facility, The Hospital for Sick Children) for technical training and analyses discussions. Models were generated with BioRender.com. G.F.W.W. is gratefully supported by a Vanier Canada Graduate Scholarship from the Canadian Institutes of Health Research (CIHR) and by an MD/PhD Studentship from the University of Toronto. J.H.B. is supported by the CIHR grant FDN-154329. A.D. is funded by grant R01GM132565 from the National Institutes of Health. G.R.V.H. is funded by grant 1R35GM119412-01 from the National Institutes of Health. S.G. is supported by the CIHR grant FDN-143202, and G.D.F. is supported by CIHR project grants PJT166010 and PJT165968.

## Author contributions

G.F.W. Walpole, S. Grinstein., G.R.V. Hammond, and G.D. Fairn conceived experiments and developed methods. G.F.W. Walpole conducted experiments and analyzed resulting data. J. Pacheco conducted photoactivation assays and analyzed resulting data with G.R.V. Hammond.

N. Chauhan, Y. M. Abbas, F. Montaño-Rendon, Z. Liu, H. Zhu, J. H. Brumell, and A. Deiters made and/or provided critical reagents. G.F.W. Walpole wrote the original draft of the manuscript. All authors reviewed and edited the manuscript.

---

**Supplementary Fig. 1, associated with Figure 1. Rapid and sustained PtdIns(3,4)P_2_ synthesis during *Salmonella* entry and maturation.**

(A) Schematic representation of the PtdIns(3,4)P_2_ biosensors based on single, double-, or triple-tandem carboxy-terminal PH domains from human TAPP1 (PLEKHA1). NES, nuclear export signal. Inset, gel electrophoresis of PCR amplicons generated by primers that house the open reading frames.

(B) Cells expressing cPHx1, cPHx2, or cPHx3 as indicated were infected for 10 min with wild-type *Salmonella* (RFP, red) prior to staining the PM (blue). Representative extended focus intensity projections (main) and a corresponding confocal section of the invasion ruffle (bottom panels) are presented.

(C) Time-lapse confocal imaging of cells (three examples in I, II, and III) expressing cPHx3 (green) during invasion by wild-type *Salmonella* (red). Bottom inset panels are expanded from the hashed box region and correspond to the minute-by-minute time series. cPHx3 is also presented using an inverted lookup table (gray) for easier visualization. See also Movies 1 and 2.

(D) As in part (C), three examples (I, II, and III) of cells expressing the PtdIns(3,4,5)P_3_ sensor aPHx2 (green and inverted gray in insets below) during invasion by wild-type *Salmonella* (red). See also Supplementary Movie 3.

(E) Sustained PtdIns(3,4)P_2_ levels during infection are SopB-dependent. The wild-type *Salmonella* dataset of Figure 1C is presented side-by-side an analysis of PM cPHx3 intensity in cells infected with an isogenic Δ*sopB* strain. Data were acquired in parallel for each bacterial strain and are statistically analyzed as in Figure 1C.

(F) Sustained Akt activation during *Salmonella* invasion is SopB-dependent. Cells serum-starved for 3 hours (to reduce basal PI3K signaling), were exposed to *Salmonella* (wild-type or Δ*sopB*) for 10 min. Extracellular bacteria were removed by extensive washing, and cells were returned to serum-free medium with gentamycin (see Methods). Lysates were collected at the indicated time-points and analyzed on parallel membranes for pAkt (S473) or pAkt (T308) prior to stripping and re-probing for total Akt or GAPDH (loading control). A series of representative immunoblots are presented from one of two independent trials. Phosphorylated residues (in parentheses) correspond to positions in Akt isoform 1, even though antibodies are not isoform specific. UI, uninfected.

**Supplementary Fig. 2, associated with Figure 2. Scarcity of PtdIns(3,4,5)P_3_ during *Salmonella* invasion or during optogenetic activation of SopB.**

(A) Schematic representation of the PtdIns(3,4,5)P_3_ biosensor NES-EGFP-aPHx2 derived from tandem ARNO pleckstrin homology domains (2G splice variant, I303E mutation). NES, nuclear export signal.

(B) Cells were exposed to invasive RFP-expressing WT or isogenic Δ*sopB Salmonella* and the PM was stained with CellMask prior to imaging. Neither recruitment of aPHx2 to CellMask-positive regions nor cytosolic depletion of the biosensor were clearly evident during invasion with either strain. Extended focus projections (main) and single confocal sections of the invasion ruffle are presented at right for each.

(C) Quantification from (B). Normalized intensity of aPHx2 in the PM of invasion sites from 3 independent trials (WT, 71 invasion sites; Δ*sopB*, 63 invasion sites). Unpaired two-tailed t-test.

(D) Normalized intensity of aPHx2 in the bulk PM during optogenetic activation of SopB wild-type (WT-464TAG) or SopB C460S (C460S-464TAG). Data are mean ± SEM (WT, 19 cells; C460S, 14 cells).

(E) Corresponding area under the curve calculations from time-lapse data in (D). Data are box (25-75^th^ percentile) and whisker (5-95^th^ percentile) plots of individual cell measurements. Compared by unpaired two-tailed t-test.

(F and G) SopB-mediated synthesis of PtdIns(3,4)P_2_ is not accompanied by a robust PtdIns(3,4,5)P_3_ response. Representative confocal sections of NES-mCherry-aPHx2 (gray) and NES-iRFP-cPHx3 (magenta) localization during optogenetic activation of WT or C460S SopB. The indicated times are prior to (t - 30 s) or after illumination with 405 nm light to photolyze hydroxycoumarin lysine in SopB. See corresponding quantification of (F) presented in Figure 2F.

**Supplementary Fig. 3, associated with Figure 3. Generation of PtdIns(3,4)P_2_ by the *Shigella* effector IpgD is resistant to PI3K inhibition.**

(A) Sequence alignment of the phosphate-binding (P-loop) sequence from SopB (*Salmonella enterica*) and IpgD (*Shigella flexneri*). Residue 439 of IpgD encodes the cysteine of the C(X)_5_R motif.

(B) Heterologous expression of EGFP-IpgD (WT or C439S) and PH-PLCδ1-mRFP1. PM was stained with CellMask prior to imaging live.

(C) Quantification of PH-PLCδ1 intensity in the PM from (B) across 3 independent trials (IpgD^WT^, 82 cells; IpgD^C439S^, 74 cells). Unpaired two-tailed student’s t-test of trial means (****, P<0.0001).

(D) Live cell imaging of heterologously-expressed EGFP-IpgD (WT or C439S) and NES-mCherry-cPHx3 following PM staining with CellMask. Cells were treated with the indicated inhibitors (PI-103, 500 nM; wortmannin, 100 nM) for 20 min prior to and throughout imaging.

(E) PM cPHx3 intensity was quantified from (D) across 3 independent trials (IpgD^WT^, 70 cells; IpgD^C439S^, 62 cells; IpgD^WT(PI-103)^, 68 cells; IpgD^WT(Wortmannin)^, 76 cells). Unpaired one-way ANOVA with Bonferroni’s multiple comparison test of trial means versus IpgD^WT^.

(F) Unprocessed immunoblots corresponding to Figure 3D and densitometry measurements in Figure 3E. The dashed red boxes indicate the regions presented in the main Figure.

**Supplementary Fig. 4, associated with Figure 4. Class II PI3K-C2α is not enriched at the site of *Salmonella invasion*.**

(A) Unprocessed immunoblots corresponding to Figure 4B and densitometry measurements in Figure 4C. The dashed red boxes indicate the regions presented in the main Figure. MW, molecular weight markers.

(B) Immunostaining (myc, red; PI3K-C2α, gray) of HeLa cells transfected with PM-targeted myc-PI3K-C2α. Non-transfected cells were overexposed in the inset to visualize endogenous staining.

(C) Confocal images of uninfected and infected HeLa cells stained for endogenous PI3K-C2α. Wild-type *Salmonella* (RFP, red; right).

**Supplementary Fig. 5, associated with Figure 8. Sequence determinants of SopB-mediated PtdIns(3,4)P_2_ generation.**

(A) Secondary structure predictions guide SopB truncation. Regions containing α-helices (blue), β-strands (magenta), or coils (black) were predicted with PSIPRED software (UCL Department of Computer Science). Plasmids beginning and ending outside major structural elements were designed to utilize natural flexible coiled regions. The position of EGFP and starting/ending residues are indicated by dashed lines.

(B and C) SopB requires amino acids 520-554 for PtdIns(3,4)P_2_ generation. The indicated constructs encoding EGFP amino-terminally fused to segments of SopB (green) were co-transfected with NES-mCherry-cPHx3 (inverted gray) and imaged live. Representative confocal sections are presented. Note that membrane-targeting of SopB is retained by both constructs, but PtdIns(3,4)P_2_ synthesis is disrupted by the deletion of 520-554.

**Supplemental Video 1, associated with** Figure 1. Live cell imaging of cPHx3 (gray, left) expressed in HeLa cells during invasion by RFP-expressing *Salmonella* (red on left). Frames were acquired at 30 sec intervals and are presented as extended focus projections of stacks of serial confocal images with 12 fps playback. cPHx3 was pseudo-coloured (right) to ease visualization, according to the inset calibration bar. See corresponding insets in Figure 1A.

**Supplemental Video 2, associated with** Figures 1 and S1. Time-lapse of fluorescence imaging of cPHx3 (gray, left) expressed during invasion by RFP-expressing *Salmonella* (red, left) delivered at high (>10) multiplicity of infection. Frames were acquired at 1 min intervals and single confocal sections are presented at 6 fps playback. cPHx3 was pseudo-coloured (right) to ease visualization according to the inset calibration bar. See additional examples in Supplementary Figure 1C.

**Supplemental Video 3, associated with** Figures 1 and S1. Time-lapse of fluorescence imaging of aPHx2 (gray) expressed during invasion by wild-type RFP-expressing *Salmonella* (red on left).

Frames were acquired at 1 min intervals, and playback is 6 fps. Merged confocal sections are presented left, and aPHx2 is pseudo-coloured right as indicated by the inset calibration bar. Additional examples are provided in Supplementary Figure 1D.

**Supplemental Video 4, associated with** Figure 2. Confocal time-lapse of NES-iRFP-cPHx3 (gray) during optogenetic activation of EGFP-SopB^WT-464-TAG^. 405-nm illumination begins at frame 15 (30 s) and remains for the duration of the video (5:29). Captures were made at 2.3 sec intervals and playback is 60 fps. See corresponding panels and quantification in Figure 2.

## References

Anderson Norris, F., Wilson, M.P., Wallis, T.S., Galyov, E.E., and Majerus, P.W. (1998). SopB, a protein required for virulence of Salmonella dublin, is an inositol phosphate phosphatase. Proc. Natl. Acad. Sci. 95, 14057–14059.

Arcaro, A., and Wymann, M.P. (1993). Wortmannin is a potent phosphatidylinositol 3-kinase inhibitor: the role of phosphatidylinositol 3,4,5-trisphosphate in neutrophil responses. Biochem. J. 296 (Pt 2, 297–301.

Balla, T. (2013). Phosphoinositides: tiny lipids with giant impact on cell regulation. Physiol. Rev. 93, 1019–1137.

Bernsen, J. (1986). Dynamic thresholding of gray-level images. In Proceedings of International Conference on Pattern Recognition, pp. 1251–1255.

Bilanges, B., Posor, Y., and Vanhaesebroeck, B. (2019). PI3K isoforms in cell signalling and vesicle trafficking. Nat. Rev. Mol. Cell Biol. 20, 515–534.

Braccini, L., Ciraolo, E., Campa, C.C., Perino, A., Longo, D.L., Tibolla, G., Pregnolato, M., Cao, Y., Tassone, B., Damilano, F., et al. (2015). PI3K-C2γ 3 is a Rab5 effector selectively controlling endosomal Akt2 activation downstream of insulin signalling. Nat. Commun. 6.

Branchu, P., Bawn, M., and Kingsley, R.A. (2018). Genome variation and molecular epidemiology of Salmonella enterica serovar Typhimurium pathovariants. Infect. Immun. 86, 1–17.

Bruno, V.M., Hannemann, S., Lara-Tejero, M., Flavell, R.A., Kleinstein, S.H., and Galán, J.E. (2009). Salmonella typhimurium type III secretion effectors stimulate innate immune responses in cultured epithelial cells. PLoS Pathog. 5.

Campbell, R.E., Tour, O., Palmer, A.E., Steinbach, P.A., Baird, G.S., Zacharias, D.A., and Tsien, R.Y. (2002). A monomeric red fluorescent protein. Proc. Natl. Acad. Sci. U. S. A. 99, 7877–7882.

Cooper, K.G., Winfree, S., Malik-Kale, P., Jolly, C., Ireland, R., Knodler, L.A., and Steele-Mortimer, O. (2011). Activation of Akt by the bacterial inositol phosphatase, SopB, is wortmannin insensitive. PLoS One 6, e22260.

Cormack, B.P., Valdivia, R.H., and Falkow, S. (1996). FACS-optimized mutants of the green fluorescent protein (GFP). Gene 173, 33–38.

Courtney, T.M., and Deiters, A. (2019). Optical control of protein phosphatase function. Nat. Commun. 10, 1–10.

Cronin, T.C., DiNitto, J.P., Czech, M.P., and Lambright, D.G. (2004). Structural determinants of phosphoinositide selectivity in splice variants of Grp1 family PH domains. EMBO J. 23, 3711–3720.

Denu, J.M., and Dixon, J.E. (1998). Protein tyrosine phosphatases: mechanisms of catalysis and regulation. Curr. Opin. Chem. Biol. 2, 633–641.

Dickson, E.J., and Hille, B. (2019). Understanding phosphoinositides: Rare, dynamic, and essential membrane phospholipids. Biochem. J. 476, 1–23.

Domin, J., Pages, F., Volinia, S., Rittenhouse, S.E., Zvelebil, M.J., Stein, R.C., and Waterfield, M.D. (1997). Cloning of a human phosphoinositide 3-kinase with a C2 domain that displays reduced sensitivity to the inhibitor wortmannin. Biochem. J. 326 (Pt 1, 139–147.

Dowler, S., Currie, R.A., Campbell, D.G., Deak, M., Kular, G., Downes, C.P., and Alessi, D.R. (2000). Identification of pleckstrin-homology-domain-containing proteins with novel phosphoinositide-binding specificities. Biochem. J. 351, 19–31.

Ebner, M., Lučić, I., Leonard, T.A., and Yudushkin, I. (2017). PI(3,4,5)P3 Engagement Restricts Akt Activity to Cellular Membranes. Mol. Cell 65, 416–431.e6.

Feng, Y., Wente, S.R., and Majerus, P.W. (2001). Overexpression of the inositol phosphatase SopB in human 293 cells stimulates cellular chloride influx and inhibits nuclear mRNA export. Proc. Natl. Acad. Sci. U. S. A. 98, 875–879.

Filonov, G.S., Piatkevich, K.D., Ting, L.M., Zhang, J., Kim, K., and Verkhusha, V. V. (2011). Bright and stable near-infrared fluorescent protein for in vivo imaging. Nat. Biotechnol. 29, 757–761.

Finn, C.E., Chong, A., Cooper, K.G., Starr, T., and Steele-Mortimer, O. (2017). A second wave of Salmonella T3SS1 activity prolongs the lifespan of infected epithelial cells. PLoS Pathog. 13, 1–28.

Folkes, A.J., Ahmadi, K., Alderton, W.K., Alix, S., Baker, S.J., Box, G., Chuckowree, I.S., Clarke, P.A., Depledge, P., Eccles, S.A., et al. (2008). The identification of 2-(1H-indazol-4-yl)-6-(4-methanesulfonyl-piperazin-1-ylmethyl)-4-morpholin-4-yl-thieno[3,2-d]pyrimidine (GDC-0941) as a potent, selective, orally bioavailable inhibitor of class I PI3 kinase for the treatment of cancer . J. Med. Chem. 51, 5522–5532.

Fruman, D.A., and Rommel, C. (2014). PI3K and cancer: Lessons, challenges and opportunities. Nat. Rev. Drug Discov. 13, 140–156.

Galán, J.E., Lara-Tejero, M., Marlovits, T.C., and Wagner, S. (2014). Bacterial type III secretion systems: specialized nanomachines for protein delivery into target cells. Annu. Rev. Microbiol. 68, 415–438.

Galyov, E.E., Wood, M.W., Rosqvist, R., Mullan, P.B., Watson, P.R., Hedges, S., and Wallis, T.S. (1997). A secreted effector protein of Salmonella dublin is translocated into eukaryotic cells and mediates inflammation and fluid secretion in infected ileal mucosa. Mol. Microbiol. 25, 903–912.

Gewinner, C., Wang, Z.C., Richardson, A., Teruya-Feldstein, J., Etemadmoghadam, D., Bowtell, D., Barretina, J., Lin, W.M., Rameh, L., Salmena, L., et al. (2009). Evidence that inositol polyphosphate 4-phosphatase type II is a tumor suppressor that inhibits PI3K signaling. Cancer Cell 16, 115–125.

Gietz, R.D., and Woods, R.A. (2002). Transformation of yeast by lithium acetate/single-stranded carrier DNA/polyethylene glycol method. Methods Enzymol. 350, 87–96.

Goulden, B.D., Pacheco, J., Dull, A., Zewe, J.P., Deiters, A., and Hammond, G.R. V (2019). A high-avidity biosensor reveals plasma membrane PI(3,4)P2 is predominantly a class I PI3K signaling product. J. Cell Biol. 218, 1066–1079.

Gregory, J.D. (1955). The Stability of N-Ethylmaleimide and its Reaction with Sulfhydryl Groups. J. Am. Chem. Soc. 77, 3922–3923.

Gulluni, F., De Santis, M.C., Margaria, J.P., Martini, M., and Hirsch, E. (2019). Class II PI3K Functions in Cell Biology and Disease. Trends Cell Biol. 29, 339–359.

Hakim, S., Bertucci, M.C., Conduit, S.E., Vuong, D.L., and Mitchell, C.A. (2012). Inositol polyphosphate phosphatases in human disease. Curr. Top. Microbiol. Immunol. 362, 247–314.

Hammond, G.R. V, Machner, M.P., and Balla, T. (2014). A novel probe for phosphatidylinositol 4-phosphate reveals multiple pools beyond the Golgi. J. Cell Biol. 205, 113–126.

Hawkins, P.T., and Stephens, L.R. (2016). Emerging evidence of signalling roles for PI(3,4)P2 in Class I and II PI3K-regulated pathways. Biochem. Soc. Trans. 44, 307–314.

Heim, R., and Tsien, R.Y. (1996). Engineering green fluorescent protein for improved brightness, longer wavelengths and fluorescence resonance energy transfer. Curr. Biol. 6, 178–182.

Herman, P.K., and Emr, S.D. (1990). Characterization of VPS34, a gene required for vacuolar protein sorting and vacuole segregation in Saccharomyces cerevisiae. Mol. Cell. Biol. 10, 6742–6754.

Hoiseth, S.K., and Stocker, B.A. (1981). Aromatic-dependent Salmonella typhimurium are non-virulent and effective as live vaccines. Nature 291, 238–239.

Horger, K.S., Estes, D.J., Capone, R., and Mayer, M. (2009). Films of agarose enable rapid formation of giant liposomes in solutions of physiologic ionic strength. J. Am. Chem. Soc. 131, 1810–1819.

Hsu, F., and Mao, Y. (2015). The structure of phosphoinositide phosphatases: Insights into substrate specificity and catalysis. Biochim. Biophys. Acta 1851, 698–710.

Hu, G.Q., Song, P.X., Chen, W., Qi, S., Yu, S.X., Du, C.T., Deng, X.M., Ouyang, H.S., and Yang, Y.J. (2017). Cirtical role for Salmonella effector SopB in regulating inflammasome activation. Mol. Immunol. 90, 280– 286.

Inoue, T., Heo, W. Do, Grimley, J.S., Wandless, T.J., and Meyer, T. (2005). An inducible translocation strategy to rapidly activate and inhibit small GTPase signaling pathways. Nat. Methods 2, 415–418.

Klarlund, J.K., Tsiaras, W., Holik, J.J., Chawla, A., and Czech, M.P. (2000). Distinct polyphosphoinositide binding selectivities for pleckstrin homology domains of GRP1-like proteins based on diglycine versus triglycine motifs. J. Biol. Chem. 275, 32816–32821.

Knight, Z.A., Gonzalez, B., Feldman, M.E., Zunder, E.R., Goldenberg, D.D., Williams, O., Loewith, R., Stokoe, D., Balla, A., Toth, B., et al. (2006). A pharmacological map of the PI3-K family defines a role for p110alpha in insulin signaling. Cell 125, 733–747.

Knodler, L.A., Finlay, B.B., and Steele-Mortimer, O. (2005). The Salmonella effector protein SopB protects epithelial cells from apoptosis by sustained activation of Akt. J. Biol. Chem. 280, 9058–9064.

Kum, W.W.S., Lo, B.C., Yu, H.B., and Finlay, B.B. (2011). Protective role of Akt2 in Salmonella enterica serovar Typhimurium-induced gastroenterocolitis. Infect. Immun. 79, 2554–2566.

LaRock, D.L., Chaudhary, A., and Miller, S.I. (2015). Salmonellae interactions with host processes. Nat. Rev. Microbiol. 13, 191–205.

Leibiger, B., Moede, T., Uhles, S., Barker, C.J., Creveaux, M., Domin, J., Berggren, P.-O., and Leibiger, I.B. (2010). Insulin-feedback via PI3K-C2alpha activated PKBalpha/Akt1 is required for glucose-stimulated insulin secretion. FASEB J. 24, 1824–1837.

Levin, R., Hammond, G.R. V, Balla, T., De Camilli, P., Fairn, G.D., and Grinstein, S. (2017). Multiphasic dynamics of phosphatidylinositol 4-phosphate during phagocytosis. Mol. Biol. Cell 28, 128–140.

Lien, E.C., Dibble, C.C., and Toker, A. (2017). PI3K signaling in cancer: beyond AKT. Curr. Opin. Cell Biol. 45, 62–71.

Liu, S.-L., Wang, Z.-G., Hu, Y., Xin, Y., Singaram, I., Gorai, S., Zhou, X., Shim, Y., Min, J.-H., Gong, L.-W., et al. (2018). Quantitative Lipid Imaging Reveals a New Signaling Function of Phosphatidylinositol-3,4-Bisphophate: Isoform- and Site-Specific Activation of Akt. Mol. Cell 71, 1092–1104.e5.

Lord, S.J., Velle, K.B., Mullins, R.D., and Fritz-Laylin, L.K. (2020). SuperPlots: Communicating reproducibility and variability in cell biology. J. Cell Biol. 219.

Luo, J., Uprety, R., Naro, Y., Chou, C., Nguyen, D.P., Chin, J.W., and Deiters, A. (2014). Genetically encoded optochemical probes for simultaneous fluorescence reporting and light activation of protein function with two-photon excitation. J. Am. Chem. Soc. 136, 15551–15558.

Maehama, T., Taylor, G.S., Slama, J.T., and Dixon, J.E. (2000). A sensitive assay for phosphoinositide phosphatases. Anal. Biochem. 279, 248–250.

Mallo, G. V., Espina, M., Smith, A.C., Terebiznik, M.R., Alemán, A., Finlay, B.B., Rameh, L.E., Grinstein, S., and Brumell, J.H. (2008). SopB promotes phosphatidylinositol 3-phosphate formation on Salmonella vacuoles by recruiting Rab5 and Vps34. J. Cell Biol. 182, 741–752.

Manna, D., Albanese, A., Park, W.S., and Cho, W. (2007). Mechanistic basis of differential cellular responses of phosphatidylinositol 3,4-bisphosphate- and phosphatidylinositol 3,4,5-trisphosphate-binding pleckstrin homology domains. J. Biol. Chem. 282, 32093–32105.

Marat, A.L., and Haucke, V. (2016). Phosphatidylinositol 3-phosphates-at the interface between cell signalling and membrane traffic. EMBO J. 35, 561–579.

Marcus, S.L., Wenk, M.R., Steele-Mortimer, O., and Finlay, B.B. (2001). A synaptojanin-homologous region of Salmonella typhimurium SigD is essential for inositol phosphatase activity and Akt activation. FEBS Lett. 494, 201–207.

Mason, D., Mallo, G. V, Terebiznik, M.R., Payrastre, B., Finlay, B.B., Brumell, J.H., Rameh, L., and Grinstein, S. (2007). Alteration of epithelial structure and function associated with PtdIns(4,5)P2 degradation by a bacterial phosphatase. J. Gen. Physiol. 129, 267–283.

McCrea, H.J., and De Camilli, P. (2009). Mutations in phosphoinositide metabolizing enzymes and human disease. Physiology (Bethesda). 24, 8–16.

Niebuhr, K., Giuriato, S., Pedron, T., Philpott, D.J., Gaits, F., Sable, J., Sheetz, M.P., Parsot, C., Sansonetti, P.J., and Payrastre, B. (2002). Conversion of PtdIns(4,5)P(2) into PtdIns(5)P by the S.flexneri effector IpgD reorganizes host cell morphology. EMBO J. 21, 5069–5078.

Norris, F.A., Atkins, R.C., and Majerus, P.W. (1997). The cDNA cloning and characterization of inositol polyphosphate 4-phosphatase type II. Evidence for conserved alternative splicing in the 4-phosphatase family. J. Biol. Chem. 272, 23859–23864.

Olivo-Marin, J.-C. (2002). Extraction of spots in biological images using multiscale products. Pattern Recognit. 35, 1989–1996.

Pendaries, C., Tronchère, H., Arbibe, L., Mounier, J., Gozani, O., Cantley, L., Fry, M.J., Gaits-Iacovoni, F., Sansonetti, P.J., and Payrastre, B. (2006). PtdIns5P activates the host cell PI3-kinase/Akt pathway during Shigella flexneri infection. EMBO J. 25, 1024–1034.

Pizarro-Cerdá, J., Kühbacher, A., and Cossart, P. (2015). Phosphoinositides and host-pathogen interactions. Biochim. Biophys. Acta 1851, 911–918.

Posor, Y., Eichhorn-Gruenig, M., Puchkov, D., Schöneberg, J., Ullrich, A., Lampe, A., Müller, R., Zarbakhsh, S., Gulluni, F., Hirsch, E., et al. (2013). Spatiotemporal control of endocytosis by phosphatidylinositol-3,4-bisphosphate. Nature 499, 233–237.

Posor, Y., Eichhorn-Grünig, M., and Haucke, V. (2015). Phosphoinositides in endocytosis. Biochim. Biophys. Acta - Mol. Cell Biol. Lipids 1851, 794–804.

Robinson, J.S., Klionsky, D.J., Banta, L.M., and Emr, S.D. (1988). Protein sorting in Saccharomyces cerevisiae: isolation of mutants defective in the delivery and processing of multiple vacuolar hydrolases. Mol. Cell. Biol. 8, 4936–4948.

Rogers, L.D., Brown, N.F., Fang, Y., Pelech, S., and Foster, L.J. (2011). Phosphoproteomic analysis of Salmonella-infected cells identifies key kinase regulators and SopB-dependent host phosphorylation events. Sci. Signal. 4, 1–14.

Sasaki, T., Takasuga, S., Sasaki, J., Kofuji, S., Eguchi, S., Yamazaki, M., and Suzuki, A. (2009). Mammalian phosphoinositide kinases and phosphatases. Prog. Lipid Res. 48, 307–343.

Scanu, T., Spaapen, R.M., Bakker, J.M., Pratap, C.B., Wu, L. en, Hofland, I., Broeks, A., Shukla, V.K., Kumar, M., Janssen, H., et al. (2015). Salmonella Manipulation of Host Signaling Pathways Provokes Cellular Transformation Associated with Gallbladder Carcinoma. Cell Host Microbe 17, 763–774.

Schindelin, J., Arganda-Carreras, I., Frise, E., Kaynig, V., Longair, M., Pietzsch, T., Preibisch, S., Rueden, C., Saalfeld, S., Schmid, B., et al. (2012). Fiji: an open-source platform for biological-image analysis. Nat. Methods 9, 676–682.

Schroeder, G.N., and Hilbi, H. (2008). Molecular pathogenesis of Shigella spp.: controlling host cell signaling, invasion, and death by type III secretion. Clin. Microbiol. Rev. 21, 134–156.

Schu, P. V, Takegawa, K., Fry, M.J., Stack, J.H., Waterfield, M.D., and Emr, S.D. (1993). Phosphatidylinositol 3-kinase encoded by yeast VPS34 gene essential for protein sorting. Science 260, 88–91.

Serunian, L.A., Haber, M.T., Fukui, T., Kim, J.W., Rhee, S.G., Lowenstein, J.M., and Cantley, L.C. (1989). Polyphosphoinositides produced by phosphatidylinositol 3-kinase are poor substrates for phospholipases C from rat liver and bovine brain. J. Biol. Chem. 264, 17809–17815.

Shaner, N.C., Campbell, R.E., Steinbach, P.A., Giepmans, B.N.G., Palmer, A.E., and Tsien, R.Y. (2004). Improved monomeric red, orange and yellow fluorescent proteins derived from Discosoma sp. red fluorescent protein. Nat. Biotechnol. 22, 1567–1572.

Smal, I., Loog, M., Niessen, W., and Meijering, E. (2010). Quantitative comparison of spot detection methods in fluorescence microscopy. IEEE Trans. Med. Imaging 29, 282–301.

Stack, J.H., DeWald, D.B., Takegawa, K., and Emr, S.D. (1995). Vesicle-mediated protein transport: regulatory interactions between the Vps15 protein kinase and the Vps34 PtdIns 3-kinase essential for protein sorting to the vacuole in yeast. J. Cell Biol. 129, 321–334.

Stauffer, T.P., Ahn, S., and Meyer, T. (1998). Receptor-induced transient reduction in plasma membrane PtdIns(4,5)P2 concentration monitored in living cells. Curr. Biol. 8, 343–346.

Steele-Mortimer, O., Meresse, S., Gorvel, J.P., Toh, B.H., and Finlay, B.B. (1999). Biogenesis of Salmonella typhimurium-containing vacuoles in epithelial cells involves interactions with the early endocytic pathway. Cell. Microbiol 1, 33–49.

Steele-Mortimer, O., Knodler, L.A., Marcus, S.L., Scheid, M.P., Goh, B., Pfeifer, C.G., Duronio, V., and Finlay, B.B. (2000). Activation of Akt/protein kinase B in epithelial cells by the Salmonella typhimurium effector sigD. J. Biol. Chem. 275, 37718–37724.

Stephens, L.R., Jackson, T.R., and Hawkins, P.T. (1993). Agonist-stimulated synthesis of phosphatidylinositol(3,4,5)-trisphosphate: a new intracellular signalling system? Biochim. Biophys. Acta 1179, 27–75.

Subach, O.M., Cranfill, P.J., Davidson, M.W., and Verkhusha, V. V. (2011). An enhanced monomeric blue fluorescent protein with the high chemical stability of the chromophore. PLoS One 6.

Tahoun, A., Mahajan, S., Paxton, E., Malterer, G., Donaldson, D.S., Wang, D., Tan, A., Gillespie, T.L., O’Shea, M., Roe, A.J., et al. (2012). Salmonella transforms follicle-associated epithelial cells into M cells to promote intestinal invasion. Cell Host Microbe 12, 645–656.

Terebiznik, M.R., Vieira, O. V, Marcus, S.L., Slade, A., Yip, C.M., Trimble, W.S., Meyer, T., Finlay, B.B., and Grinstein, S. (2002). Elimination of host cell PtdIns(4,5)P(2) by bacterial SigD promotes membrane fission during invasion by Salmonella. Nat. Cell Biol. 4, 766–773.

Teruel, M.N., and Meyer, T. (2000). Translocation and reversible localization of signaling proteins: A dynamic future for signal transduction. Cell 103, 181–184.

Thomas, C.C., Dowler, S., Deak, M., Alessi, D.R., and van Aalten, D.M. (2001). Crystal structure of the phosphatidylinositol 3,4-bisphosphate-binding pleckstrin homology (PH) domain of tandem PH-domain-containing protein 1 (TAPP1): molecular basis of lipid specificity. Biochem. J. 358, 287–294.

Tóth, J.T., Gulyás, G., Tóth, D.J., Balla, A., Hammond, G.R.V., Hunyady, L., Balla, T., and Várnai, P. (2016). BRET-monitoring of the dynamic changes of inositol lipid pools in living cells reveals a PKC-dependent PtdIns4P increase upon EGF and M3 receptor activation. Biochim. Biophys. Acta - Mol. Cell Biol. Lipids 1861, 177–187.

Valdivia, R.H., Hromockyj, A.E., Monack, D., Ramakrishnan, L., and Falkow, S. (1996). Applications for green fluorescent protein (GFP) in the study of host-pathogen interactions. Gene 173, 47–52.

Vanhaesebroeck, B., Guillermet-Guibert, J., Graupera, M., and Bilanges, B. (2010). The emerging mechanisms of isoform-specific PI3K signalling. Nat. Rev. Mol. Cell Biol. 11, 329–341.

Vanhaesebroeck, B., Stephens, L., and Hawkins, P. (2012). PI3K signalling: The path to discovery and understanding. Nat. Rev. Mol. Cell Biol. 13, 195–203.

Varnai, P., Thyagarajan, B., Rohacs, T., and Balla, T. (2006). Rapidly inducible changes in phosphatidylinositol 4,5-bisphosphate levels influence multiple regulatory functions of the lipid in intact living cells. J. Cell Biol. 175, 377–382.

Várnai, P., and Balla, T. (1998). Visualization of phosphoinositides that bind pleckstrin homology domains: Calcium- and agonist-induced dynamic changes and relationship to myo-[3H]inositol-labeled phosphoinositide pools. J. Cell Biol. 143, 501–510.

Venkateswarlu, K., Oatey, P.B., Tavaré, J.M., and Cullen, P.J. (1998). Insulin-dependent translocation of ARNO to the plasma membrane of adipocytes requires phosphatidylinositol 3-kinase. Curr. Biol. 8, 463– 466.

Virbasius, J. V., Guilherme, A., and Czech, M.P. (1996). Mouse p170 is a novel phosphatidylinositol 3-kinase containing a C2 domain. J. Biol. Chem. 271, 13304–13307.

Walpole, G.F.W., and Grinstein, S. (2020). Endocytosis and the internalization of pathogenic organisms: focus on phosphoinositides. F1000Research 9, 368.

Wennström, S., and Downward, J. (1999). Role of Phosphoinositide 3-Kinase in Activation of Ras and Mitogen-Activated Protein Kinase by Epidermal Growth Factor. Mol. Cell. Biol. 19, 4279–4288.

Willars, G.B., Nahorski, S.R., and Challiss, R.A.J. (1998). Differential regulation of muscarinic acetylcholine receptor-sensitive polyphosphoinositide pools and consequences for signaling in human neuroblastoma cells. J. Biol. Chem. 273, 5037–5046.

Won, D.H., Inoue, T., Wei, S.P., Man, L.K., Byung, O.P., Wandless, T.J., and Meyer, T. (2006). PI(3,4,5)P3 and PI(4,5)P2 lipids target proteins with polybasic clusters to the plasma membrane. Science (80-.). 314, 1458–1461.

Zhang, K., Riba, A., Nietschke, M., Torow, N., Repnik, U., Pütz, A., Fulde, M., Dupont, A., Hensel, M., and Hornef, M. (2018). Minimal SPI1-T3SS effector requirement for Salmonella enterocyte invasion and intracellular proliferation in vivo. PLoS Pathog. 14, e1006925.

Zhang, S., Santos, R.L., Tsolis, R.M., Stender, S., Hardt, W.-D., Bäumler, A.J., and Adams, L.G. (2002). The Salmonella enterica serotype typhimurium effector proteins SipA, SopA, SopB, SopD, and SopE2 act in concert to induce diarrhea in calves. Infect. Immun. 70, 3843–3855.

Zhu, L., Jorgensen, J.R., Li, M., Chuang, Y.S., and Emr, S.D. (2017). ESCRTS function directly on the lysosome membrane to downregulate ubiquitinated lysosomal membrane proteins. Elife 6, 1–20.

